# Myogenesis modelled by human pluripotent stem cells uncovers Duchenne muscular dystrophy phenotypes prior to skeletal muscle commitment

**DOI:** 10.1101/720920

**Authors:** Virginie Mournetas, Emmanuelle Massouridès, Jean-Baptiste Dupont, Etienne Kornobis, Hélène Polvèche, Margot Jarrige, Maxime R. F. Gosselin, Antigoni Manousopoulou, Spiros D. Garbis, Dariusz C. Górecki, Christian Pinset

## Abstract

Duchenne muscular dystrophy (DMD) causes severe disability of children and death of young men, with an incidence of approximately 1/5,000 male births. Symptoms appear in early childhood, with a diagnosis made around 4 years old, a time where the amount of muscle damage is already significant, preventing early therapeutic interventions that could be more efficient at halting disease progression. In the meantime, the precise moment at which disease phenotypes arise – even asymptomatically – is still unknown. Thus, there is a critical need to better define DMD onset as well as its first manifestations, which could help identify early disease biomarkers and novel therapeutic targets.

In this study, we have used human induced pluripotent stem cells (hiPSCs) from DMD patients to model skeletal myogenesis, and compared their differentiation dynamics to that of healthy control cells by a comprehensive multi-omic analysis. Transcriptome and miRnome comparisons combined with protein analyses at 7 time points demonstrated that hiPSC differentiation 1) mimics described DMD phenotypes at the differentiation endpoint; and 2) homogeneously and robustly recapitulates key developmental steps - mesoderm, somite, skeletal muscle - which offers the possibility to explore dystrophin functions and find earlier disease biomarkers.

Starting at the somite stage, mitochondrial gene dysregulations escalate during differentiation. We also describe fibrosis as an intrinsic feature of skeletal muscle cells that starts early during myogenesis. In sum, our data strongly argue for an early developmental manifestation of DMD whose onset is triggered before the entry into the skeletal muscle compartment, data leading to a necessary reconsideration of dystrophin functions during muscle development.

## INTRODUCTION

Duchenne muscular dystrophy (DMD) is a rare genetic disease, but it is the most common form of myopathy affecting approximately one in 5,000 male births and very rarely female. In this recessive X-linked monogenic disorder, mutations in the *DMD* gene lead to the loss of functional dystrophin protein, resulting in a progressive - yet severe - muscle wasting phenotype (1). In patients, symptoms usually appear in early childhood (2-5 years old) and worsen with age, imposing the use of wheelchair before 15 and leading to premature death by cardiac and/or respiratory failure(s) mostly around 30 years of age (2).

At the age of diagnosis (around 4 years old), muscles of DMD patients have already suffered from the pathology (3,4). Several reviews pointed out the limitations of current disease biomarkers, which fail to detect the development of DMD specifically and at an early age (5,6). Meanwhile, no treatment is available to stop this degenerative disease yet. Developing therapies aim at restoring the expression of dystrophin in muscle cells but, so far, the level stays too low to be beneficial to patients (7). The absence of both reliable biomarkers and effective therapies stress the need of better defining the first steps of DMD in humans to be able to increase diagnosis sensitivity and, therefore, improve patient management by accelerating their access to better healthcare as well as develop alternative therapeutic approaches by finding targets that compensate the lack of dystrophin and complement current attempts at restoring its expression (8).

In 2007, a seminal publication reported that the gene expression profile of muscles from asymptomatic DMD children younger than 2 years old is already distinguishable from healthy muscles, suggesting that DMD molecular dysregulations appear before disease symptomatic manifestations (4). Evidence obtained in multiple animal models, such as neonatal *GRMD* dogs (9), DMD zebrafish (10) and *mdx* mouse embryos (11), as well as in human foetuses (12–14) even suggest that DMD starts before birth, during prenatal development. Our team recently identified the embryonic dystrophin isoform Dp412e expressed in early mesoderm-committed cells (15), another indication that DMD can start *in utero*. Further exploring DMD onset in human foetuses is extremely challenging for obvious ethical and practical reasons. A way to overcome these issues is to develop a human DMD model *in vitro*, recapitulating embryonic development from human pluripotent stem cells to skeletal muscle lineage.

To our knowledge, none of the existing human DMD *in vitro* models, either based on tissue-derived myoblasts (16) or on the differentiation of induced pluripotent stem cells (17–21), have been used for studying DMD during the ontogeny of the skeletal muscle lineage. Moreover, original protocols for *in vitro* myogenesis from human pluripotent stem cells (reviewed in (22)) use transgene overexpression or/and cell sorting procedures, and thereby, miss the steps preceding skeletal muscle commitment, *e.g.* paraxial mesoderm and myotome. Novel protocols have recently used transgene-free directed differentiation to recapitulate human embryonic development in a dish, giving theoretical access to the developmental steps (19,23–25).

Using one of these protocols (23), we compared the myogenic differentiation dynamics of healthy and DMD hiPSCs using a multi-omic approach to identify early disease manifestations *in vitro*. DMD cells showed marked transcriptome dysregulations from day 10, before the detection of skeletal muscle regulatory factors at day 17. Specifically, we identified the dysregulation of mitochondrial genes as one of the earliest detectable phenotypes. These alterations escalated over the course of muscle specification. In addition, we showed an early induction of Sonic hedgehog signalling pathway, followed by collagens as well as fibrosis-related genes suggesting the existence of an intrinsic fibrotic process solely driven by DMD muscle cells. Overall, our data highlight that human pluripotent stem cells are a suitable cell model to study the ontogeny of skeletal muscle lineage in both healthy and disease conditions. In the context of DMD, they strongly argue for the existence of early disease manifestations during somite development.

## RESULTS

To establish the early/developmental impact of *DMD* gene mutations, human induced pluripotent stem cells (hiPSCs) from three DMD patients and three healthy individuals were generated as described previously (15). These cells were subjected to a standardised differentiation protocol without utilisation of feeder cells, cell sorting or gene overexpression resulting in elongated and plurinucleated myotubes within 25 days (23), with an amplification fold of 2918 ± 480 (mean ± SEM). Skeletal muscle progenitor cells after 10 and 17 days of differentiation could be cryopreserved (Figure S1A). Whole transcriptome and miRnome profiles were compared at 7 differentiation time points (tissue-derived myoblasts and myotubes, as well as hiPSC-derived cells at days 0, 3, 10, 17 and 25) and complemented by TMT proteomics and Western blot analyses (Table S1).

### DMD is initiated prior to the expression of skeletal muscle markers

First, the expression profile of the *DMD* variants was studied by RT-qPCR in healthy and DMD hiPSCs during the differentiation process described in Figure S1A. The *Dp427m* variant, which is normally observed in muscle cells (26), appeared from day 3 and was increased at day 17, in contrast with *Dp412e* – the embryonic variant of dystrophin present in mesoderm cells (15) – which was expressed from day 0, increased at differentiation day 3 and disappeared from day 10. Therefore, the expression of the *DMD* locus is initiated in the very first steps of the differentiation protocol, well before the entry into the skeletal muscle lineage. The ubiquitous variant *Dp71-40* was detected at every time points, in contrast with *Dp116* (Schwann cell variant (27)), *Dp140* (kidney and foetal brain variant (28)) *Dp427p1p2* (Purkinje cell variant (29)), and *Dp427c* which were either undetected or expressed at very low levels (Figure S1B). Interestingly, *Dp260* (retinal variant (30)) followed a similar expression pattern than *Dp427m*.

A strong correlation in the transcriptomic data was observed by mRNA-seq and miRNA-seq between samples collected at an individual time point, as opposed to samples from two distinct time points. In addition, the correlation coefficient between samples taken at two successive time points increased as differentiation progressed (Figure 1A). Differential expression analysis in healthy controls between two successive collection days (days 3/0, days 10/3, days 17/10, days 25/17) showed that the proportion of regulated genes decreased from 26 % to 18 % of the whole transcriptome through the course of differentiation (8080 to 5320 mRNAs, adjusted p-value ≤ 0.01, Figure S2A). These observations demonstrate the robustness of the differentiation protocol and are in agreement with an early specialisation and a later refinement of the transcriptome as cells quickly exit pluripotency and become progressively restricted to the skeletal muscle lineage.

**Figure 1.**
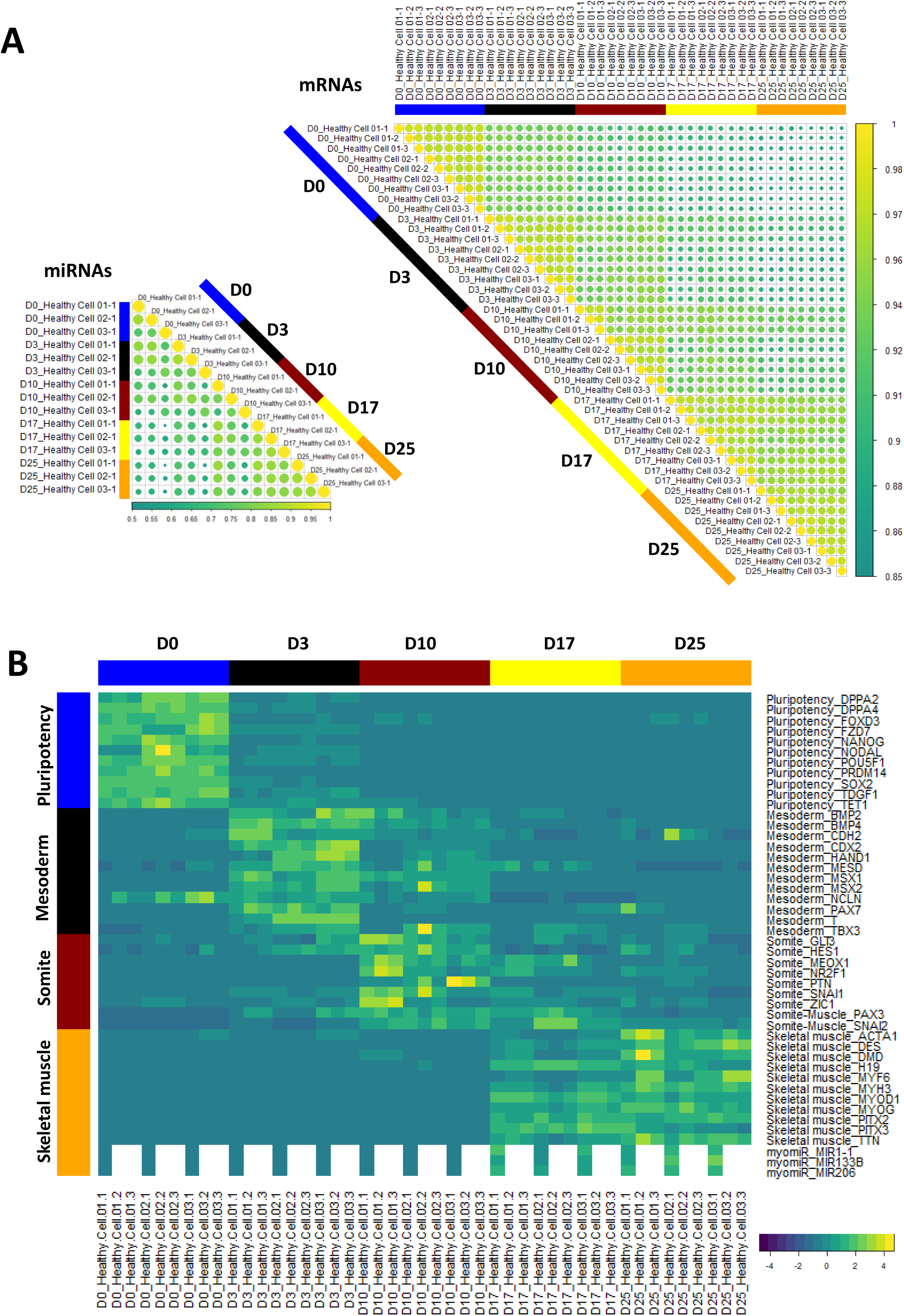
Differentiation dynamics of hiPSCs (D0) into MyoT (D25) in healthy cells at the transcriptomic level. **A)** Spearman correlation matrix of transcriptomes (mRNAs, right) and miRnomes (miRNAs, left). Yellow dots indicate a stronger correlation. **B)** Gene expression heatmap of selected differentiation markers. (D: day; hiPSC: human induced pluripotent stem cell; MyoT: myotube).

To characterise the developmental stages achieved by the cells, the expression of lineage-specific markers (both mRNAs and miRNAs) was determined at each time point, together with gene ontology enrichment analyses (Figure 1B-2A, Figure S2B-C, Table S2).

**Figure 2.**
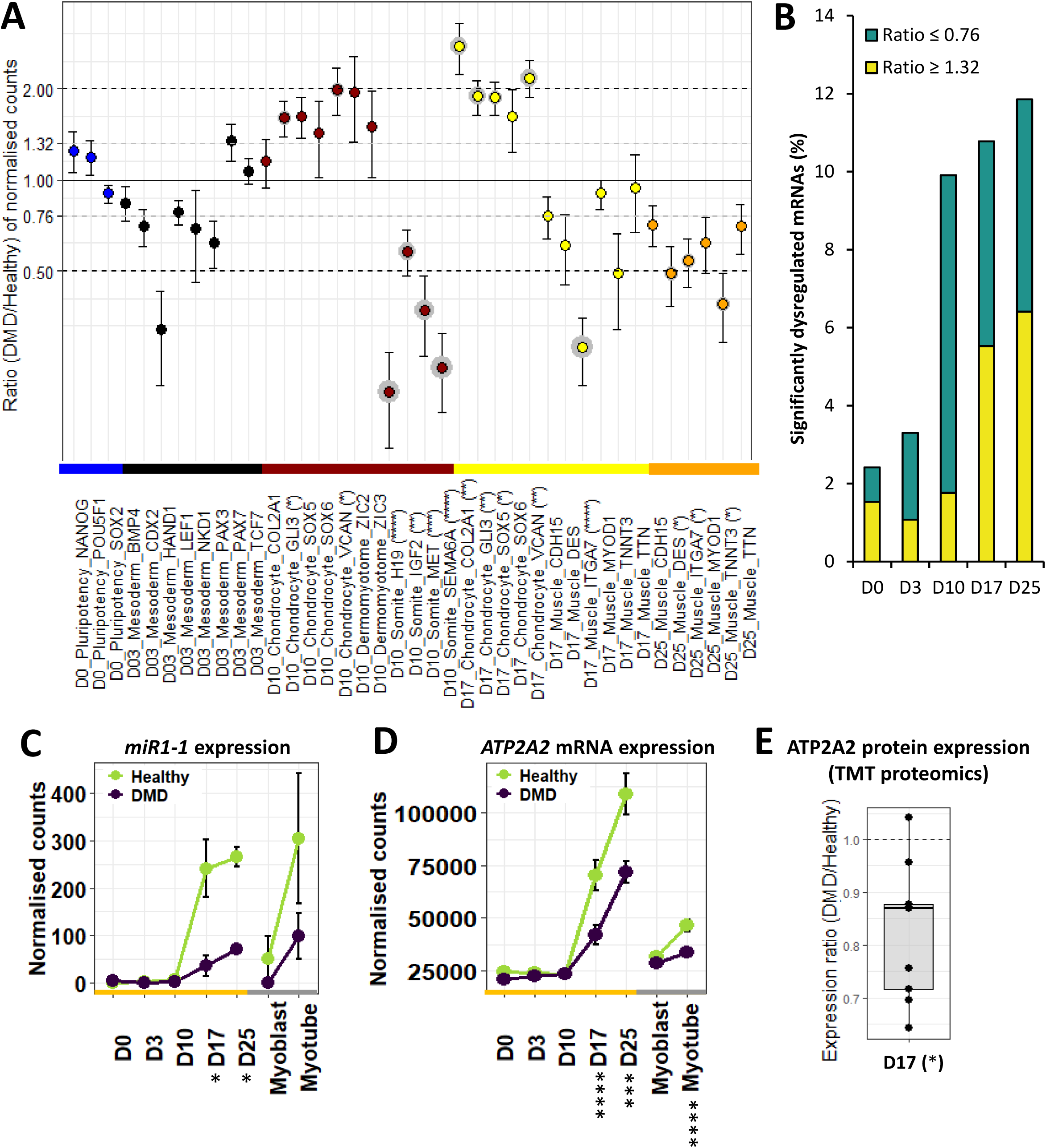
Differentiation dynamics of hiPSCs (D0) into MyoT (D25) in DMD cells. **A)** Dotplot of DMD/healthy expression ratios of selected differentiation markers. Statistical differences are indicated in brackets after gene names, and grey circles around the corresponding dots. **B)** Proportions of significantly dysregulated mRNAs (adjusted p-value ≤ 0.05) in DMD cells at each time points. Expression of **C)** *MIR1-1* and **D)** *ATP2A2* mRNA during differentiation. **E)** ATP2A2 protein level at D17. (*adjusted p-value ≤ 0.05, **adjusted p-value ≤ 0.01, ***adjusted p-value ≤ 0.001, ****adjusted p-value ≤ 0.0001; D: day; hiPSC: human induced pluripotent stem cell; MyoT: myotube).

Pluripotency was similarly maintained in healthy and DMD cells at day 0 (Figure 2A, Table S2), as already shown by our group (15). At day 3, cells lost pluripotency and became paraxial mesoderm cells expressing marker genes such as *PAX3* and *PAX7* (11) (Figure 2A, Table S2). Importantly, markers of lateral plate (*e.g. GATA4* (31)) and intermediate mesoderm (*e.g. PAX8* (32)) were not upregulated at this stage (Table S2). Similarly, earlier markers of primitive streak (*e.g. TBX6* (33)), mesendoderm (*e.g. MIXL1* (34)), as well as markers of the other germ layers, endoderm (*e.g. SOX17* (35)) and ectoderm (*e.g. SOX2* (36)) were either not expressed, greatly downregulated or expressed at very low levels (Table S2), suggesting cell homogeneity in the differentiation process.

At that early time point, DMD-associated gene dysregulation represented less than 3 % of the entire transcriptome (adjusted p-value ≤ 0.05, Figure 2B) but already contained genes important for development (*e.g. MEIS2* (37)) and muscle formation (*e.g. ACTA1* (38)). However, mesoderm markers were not significantly dysregulated, attesting that mesoderm commitment was mostly unimpaired (Figure 2A, Table S2). No increase in the expression of primitive streak, mesendoderm, endoderm or ectoderm markers was detected, suggesting no differences in the differentiation process of DMD cells at that stage (Table S2).

In contrast, a sharp increase in the proportion of dysregulated genes appeared at day 10, mostly including gene downregulations (DMD/Healthy expression ratio ≤ 0.76, adjusted p-value ≤ 0.05). This concerned almost 10 % of the transcriptome at day 10 (against 3 % at day 3) and remained stable from 10 to 12 % (1226 mRNAs) until day 25 (Figure 2B). At day 10, healthy cells expressed genes typically observed during somitogenesis, such as *PAX3* (39) *NR2F2* (40), *PTN* (41), *MET* (42), *H19* and *IGF2* (43) (Table S2). More precisely, their transcriptome exhibits a mixed profile between dermomyotome (expression of *GLI3* (44) and *GAS1* (45) but not *ZIC3* (46)) and myotome (expression of *MET* (47) and *EPHA4* (48) but not *LBX1* (49)) (Table S2). Neither markers of presomitic mesoderm cells (*e.g. FGF8* (50)) and neural plate cells (*FOXD3* (51)), nor markers of sclerotome (*e.g. PAX1* (52)) and dermatome (*e.g. EGFL6* (53)) were upregulated (Table S2) in both healthy and DMD cells. In the meantime, several somite markers were downregulated, including *H19, IGF2, MET* and *SEMA6A* (54) (validated at the protein level for SEMA6A, Figure 2A-S3A, Table S2), while a slight upregulation of chondrocyte markers was highlighted and confirmed at the protein level for GLI3 (Figure S3B), together with a significant enrichment of the gene ontology term ‘nervous system development’, suggesting potential lineage bifurcations at day 10 (Figure 2A-S2C, Table S2).

The study of differentiation dynamics presented above highlights that mesoderm commitment is not impaired by the absence of dystrophin, and shows that DMD onset takes place at the somite cell stage, before the expression of the skeletal muscle program and especially before the upregulation of *Dp427m* expression.

### DMD skeletal muscle progenitor cells exhibit specific muscle gene dysregulations

Healthy and DMD cells were in the skeletal muscle compartment at day 17, as evidenced by the expression of multiple lineage-specific genes, such as transcription factors (*e.g. MYOD1* (55)), cell surface markers (*e.g. CDH15* (56)), sarcomere genes (*e.g. TNNC2* (57)), dystrophin-associated protein complex (DAPC) genes (*e.g. SGCA* (58)), Calcium homeostasis genes (*e.g. RYR1* (59)) and muscle-specific miRNAs (myomiR, *e.g. MIR1-1* (60)). This was also observed at the protein level for CDH15, TNNC2 and RYR1 (Figure 1B, Table S2). They both showed an embryonic/foetal phenotype characterised by *ERBB3* expression, in contrast with tissue-derived myoblasts that expressed *NGFR* (21). Here again, alternative cell lineages were absent or greatly downregulated, such as tenocytes (*e.g. MKX* (61)), chondrocytes (*e.g. SOX5* (62)), osteoblasts (*e.g. SPP1* (63)) or nephron progenitors (*e.g. SALL1* (64)) (Table S2).

Interestingly, DMD cells did not show a significant dysregulation of skeletal muscle transcription factors (Table S2). However, several myomiRs were found downregulated (*e.g. MIR1-1*, Figure 2C), together with genes related to calcium homeostasis (*e.g. ATP2A2* (65), at both mRNA and protein level, Figure 2D-E) as well as members of the DAPC (*e.g. SNTA1* (66)) (Table S2). Concerning cell lineages, there was no visible difference when compared to healthy controls, except an upregulation of markers associated with chondrocytes, which was confirmed at the protein level for GLI3 (Figure S3C), and a significant enrichment of the gene ontology term ‘nervous system development’ previously seen at day 10, together with ‘kidney development’ and ‘ossification’ (Figure 2A-S2C, Table S2).

DMD-specific dysregulations were further queried at the protein level using TMT proteomics. 3826 proteins were detected in the 6 processed samples (3 healthy and 3 DMD, Table S3). Among these list, 185 proteins (139 + 46) were found significantly dysregulated in DMD and 375 (329 + 46) of the corresponding mRNAs were previously detected dysregulated in the RNA-seq analysis, the overlap between protein and mRNA identified dysregulations being 46 (|log2FoldChange| ≥ 0.4 and adjusted p-value ≤ 0.05, Figure S3D-E, Table S4). Moreover, among the total of 514 genes represented in Figure S3F, 98 were dysregulated alike in both datasets (56 upregulated + 42 downregulated) against 13 (12 + 1) in the opposite direction (|log2FoldChange| ≥ 0.4, Figure S3F, Table S4) resulting in a Spearman correlation of r = 0.49 and p-value < 0.0001. In this mRNA/protein comparison, the mRNA experiment was more sensitive than protein experiment and could also be considered as a good proxy for proteins.

To better characterise the most direct consequences of the loss of *DMD* in muscle cells, *DMD* expression was knocked-down at day 17 by transient exon skipping using a specific phosphorodiamidate morpholino oligomer targeting *DMD* exon 7 (PMO7) in a healthy hiPSC line. Treatment with PMO7 resulted in significant exon skipping which was correlated with reduced *DMD* expression up to 94% (Spearman r = −0.88, analysed pairs = 59, p-value < 0.0001, Figure S4A) and reduced dystrophin protein levels (up to 81%, Figure S4B). In parallel, the expression of specific transcripts was measured by RT-qPCR the 3 following days (Figure S4A): transcripts coding for *MYH3, MYOG* and *SGCA* were significantly downregulated after PMO7 treatment (gene group 1), while transcripts coding for *DES* and *ITGA7* were not affected (gene group 2).

Therefore, DMD cells efficiently enter the skeletal muscle compartment at day 17, but exhibit dysregulations in several features typically associated with dystrophic muscles, which could be a consequence of the early manifestations of DMD detected at day 10. Some of these identified dysregulations were mimicked by transient *DMD* knockdown.

### hiPSC differentiation leads to embryonic/foetal myotubes that reproduce DMD phenotypes

As previously described (23), both healthy and DMD hiPSC-derived myotubes (day 25) were able to twitch spontaneously in culture, and fluorescent staining of nuclei and α-actinin confirmed cell fusion and the formation of striation patterns typical of muscle fibres *in vivo* (Figure 3A). Western blot analyses on protein extracts from DMD cells confirmed that dystrophin was either undetectable or slightly expressed (Figure 3B), as in the corresponding patient muscle biopsies (data not shown).

**Figure 3.**
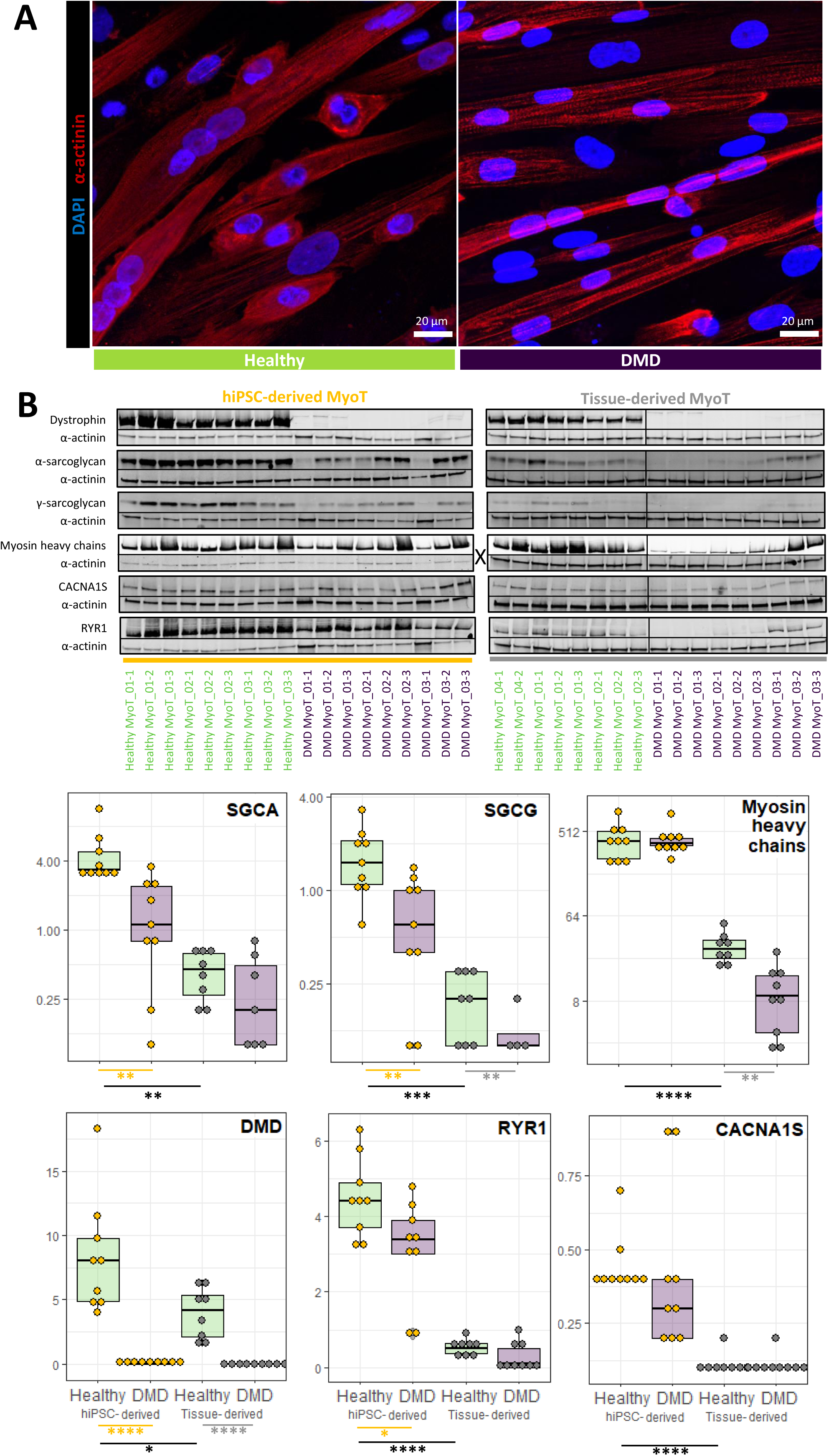
Comparison of healthy and DMD MyoT from hiPSCs and tissues at the protein level. **A)** hiPSC-derived MyoT immunolabelling of α-actinin (red) and nuclei (DAPI, blue) in healthy (left) and DMD cells (right). **B)** Representative Western blots and related quantifications of DMD, SGCA, SGCG, myosin heavy chains, CACNA1S and RYR1 from protein extracts in healthy and DMD hiPSC-derived and tissue-derived MyoT (X: 0.25 µg of total protein was used in hiPSC-derived MyoT instead of 7µg in tissue-derived MyoT - *p-value ≤ 0.05, **p-value ≤ 0.01, ***p-value ≤ 0.001, ****p-value ≤ 0.0001). (hiPSC: human induced pluripotent stem cell; MyoT: myotube).

We selected representative mRNAs and miRNAs and showed that both hiPSC-derived and tissue-derived myotubes have exited the cell cycle and upregulated genes expressed in skeletal muscles (Figure S5A, Figure 4A, Table S2). This included skeletal muscle myomiRs (*MIR1-1, MIR133* and *MIR206* (67,68)), transcription factors involved in skeletal myogenesis including those of the muscle regulatory factor (MRF) family (*e.g. MYOD1* (55), *MYOG* (69)), specific muscle cell surface markers (*e.g. CDH15* (56), *ITGA7* (70)) as well as genes involved in the formation of the DAPC (*e.g. SGCA* (58), *DTNA* (71)), sarcomeres (*e.g. TNNC2* (57), *TNNT3* (72)), myofibril organisation (*e.g. UNC45B* (73), *NACA* (74)) and the triggering of excitation-contraction coupling at the neuromuscular junction (NMJ, *e.g. MUSK* (75), *DOK7* (76)) (Figure 4A, Table S2).

**Figure 4.**
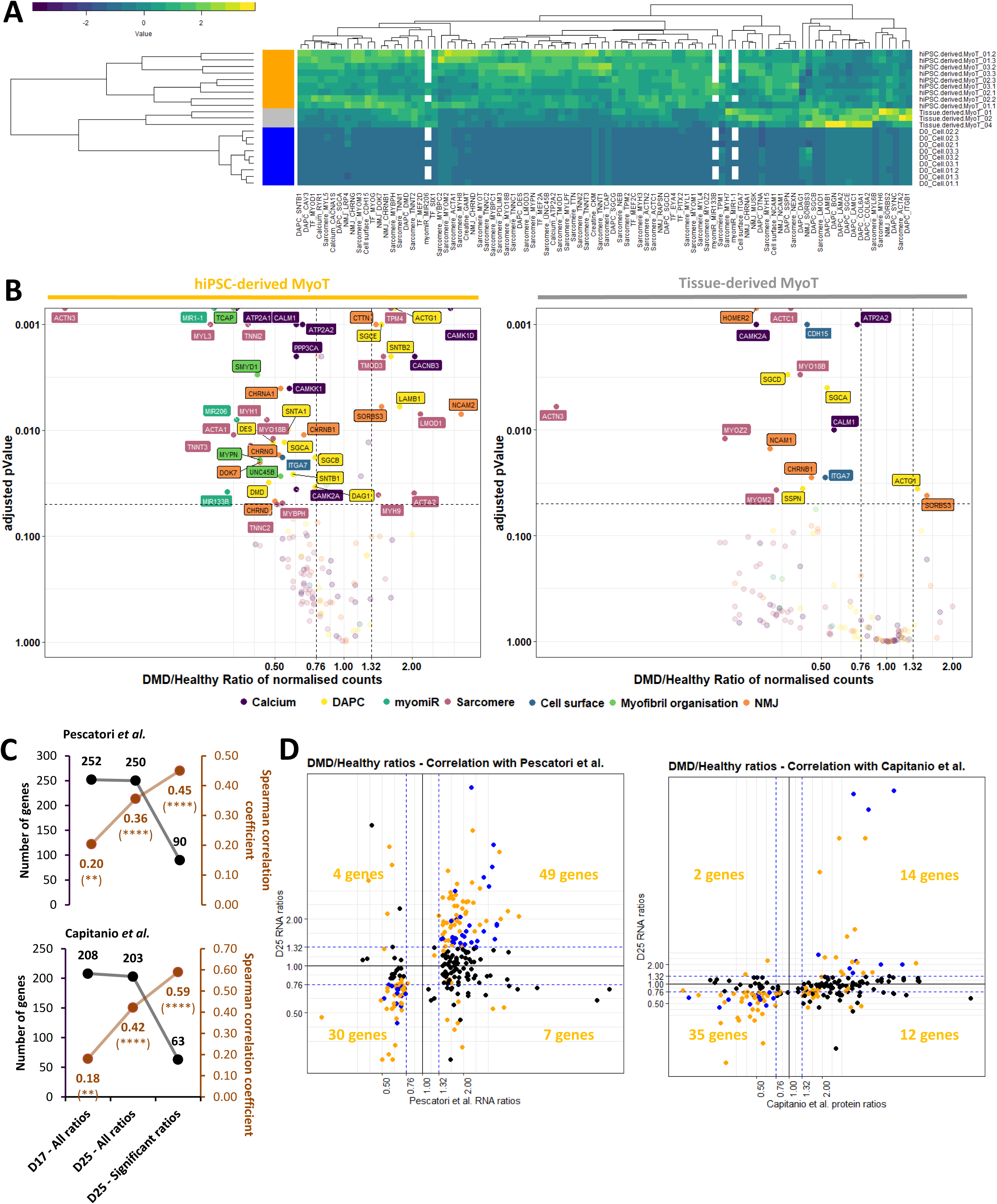
Manifestation of the DMD phenotype in the transcriptome and miRnome of myotubes derived from hiPSCs and tissues. **A)** Hierarchical clustering and heatmap in healthy hiPSCs (D0), hiPSC-derived MyoT and tissue-derived MyoT with selected skeletal muscle transcripts and miRNAs. **B)** Volcano plots of dysregulated mRNAs/miRNAs in hiPSC-derived MyoT (left) and tissue-derived MyoT (right) – vertical grey dashed lines represent DMD/Healthy ratio thresholds at 0.76 or 1.32 - the horizontal grey dashed line represents the adjusted p-value threshold at 0.05. Comparisons of DMD/Healthy expression ratios at D17 and D25 with published omics data from muscle biopsies (4,97) : **C)** number of genes in black and Spearman correlation coefficients in brown found in common with Pescatori *et al.*’s mRNA data (top) and Capitanio *et al.*’s protein data (bottom) as well as **D)** correlation graphs of the D25 data compared with Pescatori *et al.* mRNA data (left) and Capitanio *et al.* protein data (right). Genes with |log2FoldChange| ≥ 0.4 are in blue if adjusted p-value ≥ 0.05 and yellow if adjusted p-value ≤ 0.05. (DAPC: dystrophin-associated protein complex; hiPSC: human induced pluripotent stem cell; MyoT: myotube; NMJ: neuromuscular junction; TF: transcription factor MyoT - *p-value ≤ 0.05, **p-value ≤ 0.01, ***p-value ≤ 0.001, ****p-value ≤ 0.0001).

Even though global analysis showed that hiPSC-derived myotubes were similar to their tissue-derived counterparts in term of lineage commitment, they displayed an embryonic/foetal phenotype – as suggested in progenitors at day 17. This can be illustrated by the expression of the embryonic/foetal myosin heavy/light chains *MYH3* (77), *MYH8*(78), *MYL4* (79) and *MYL5* (80) but not the postnatal transcripts *MYH1* and *MYH2* (81), which were detected in tissue-derived myotubes. Myotubes derived from hiPSCs had also higher levels of *IGF2*, which is downregulated at birth (82), and expressed *DLK1*, which is known to be extinct in adult muscles (83) (Figure S5B).

Despite the embryonic/foetal phenotype, hiPSC-derived myotubes showed evidence of terminal differentiation and cellular maturation. First, their total level of myosin heavy chain proteins was significantly higher than in tissue-derived myotubes, as confirmed by Western blotting (Figure 3B). RNAs and proteins involved in DAPC formation (*e.g.* DMD, SGCA (58) and SGCG (84)), as well as in excitation-contraction coupling (*e.g.* RYR1 (59) and CACNA1S / CAV1.1 (85)) were also present at higher levels (Figure 3B-4A). Finally, higher expression of skeletal muscle transcription factors (*e.g. MEF2C* (86)), and of multiple genes involved in muscle contraction (*e.g. TNNT3* (72)), NMJ formation (*e.g. RAPSN* (87)), and creatine metabolism (*e.g. CKM* (88)) indicates that hiPSC-derived cells expressed features of fully differentiated muscle cells (Figure 4A). Similar to previous time points, day 25 cells were negative for markers of alternative muscle lineages, *i.e.* cardiac (*MIR208a* (89), *MYL7* (90) and *RYR2* (91)) and smooth muscle cells (*MYH11* (92), *CNN1* (93) and *CHRNA3/B2/B4* (94))(Table S2).

In DMD cells, unbiased mRNA-seq analysis highlighted striking transcriptome dysregulations with 3,578 differentially expressed genes in hiPSC-derived myotubes including well-known muscle genes. There was a global trend towards downregulation of muscle transcription factors, which was only significant for *MEF2A* and *MEF2D* in hiPSC-derived myotubes and *EYA4* and *MYOD1* in tissue-derived myotubes (Figure S5C). In addition, myomiRs previously associated with muscle dystrophy (dystromiRs, *e.g. MIR1-1* (60), Figure 2C) were found downregulated (Table S2). Similarly, a global downregulation phenotype was observed in both tissue- and hiPSC-derived DMD myotubes, and concerned multiple mRNAs and/or proteins associated with known disease phenotypes, such as cell surface markers (*e.g. ITGA7* (70)), DAPC organisation (*e.g.* both SGCA mRNA and protein (58) as well as SGCG protein (84)), myofibril organisation (*e.g. UNC45B* (73)), sarcomere formation (*e.g. MYO18B* (95)), NMJ function (*e.g. CHRNB1* (96)) and calcium homeostasis (*e.g. ATP2A2* mRNA (65) and RYR1 protein (59)) (Figure 3B for protein data, 4B for transcript data, S2C for enrichment data).

Then we compared the DMD/Healthy expression ratios at day 25 with two sets of published omics data from healthy and DMD muscle biopsies: one obtained at the mRNA level in pre-symptomatic DMD patients younger than 2 years old (4) and another at the protein level in patients aged from 9 months to 8 years old (97). Both datasets were closer to day 25 cells (hiPSC-derived myotubes) than day 17 cells as expected. Our hiPSC-derived myotubes expressed 250 of the 261 dysregulated genes and 203 of the 226 dysregulated proteins found in these respective studies (Spearman correlations of r = 0.36 and r = 0.42, p-value < 0.0001, Figure 4C, Table S4). Among these, respectively 90 and 63 genes were also significantly dysregulated in our dataset (|log2FoldChange| ≥ 0.4, adjusted p-value ≤ 0.05): 88% (79 / 90 genes) of the identified genes from the mRNA dataset and 78% (49 / 63 genes) of the identified genes from the protein dataset were dysregulated in the same direction, resulting in Spearman correlation of r = 0.45 and r = 0.59 respectively (p-value ≤ 0.0001, Figure 4C-D, Table S4).

Altogether, these data indicate that hiPSC-derived myotubes recapitulate a full skeletal muscle differentiation program, and exhibit an embryonic/foetal phenotype. Despite that, it shows that DMD phenotypes are already detectable at the transcriptional level and correlated with those found in human patients. This validates the relevance of this cell system to model the DMD pathology.

### Markers of fibrosis are intrinsic to DMD hiPSC-derived myotubes

As presented above, the upregulation of chondrocyte markers in DMD cells, although already present at day 10, became significant from day 17 (Figure 2A, Table S2). It was accompanied by the upregulations of the Sonic hedgehog (SHH) signalling pathway and of multiple collagens (Figure 5A, Table S2). Genes encoding the *P4H* collagen synthases, were not dysregulated while *RRBP1* (that stimulates collagen synthesis (98)) together with *PLOD1* and *PLOD2* (that stabilise collagens (99,100)) were significantly upregulated. Moreover, *SETD7*, a gene known for activating collagenases (101), was significantly downregulated.

**Figure 5.**
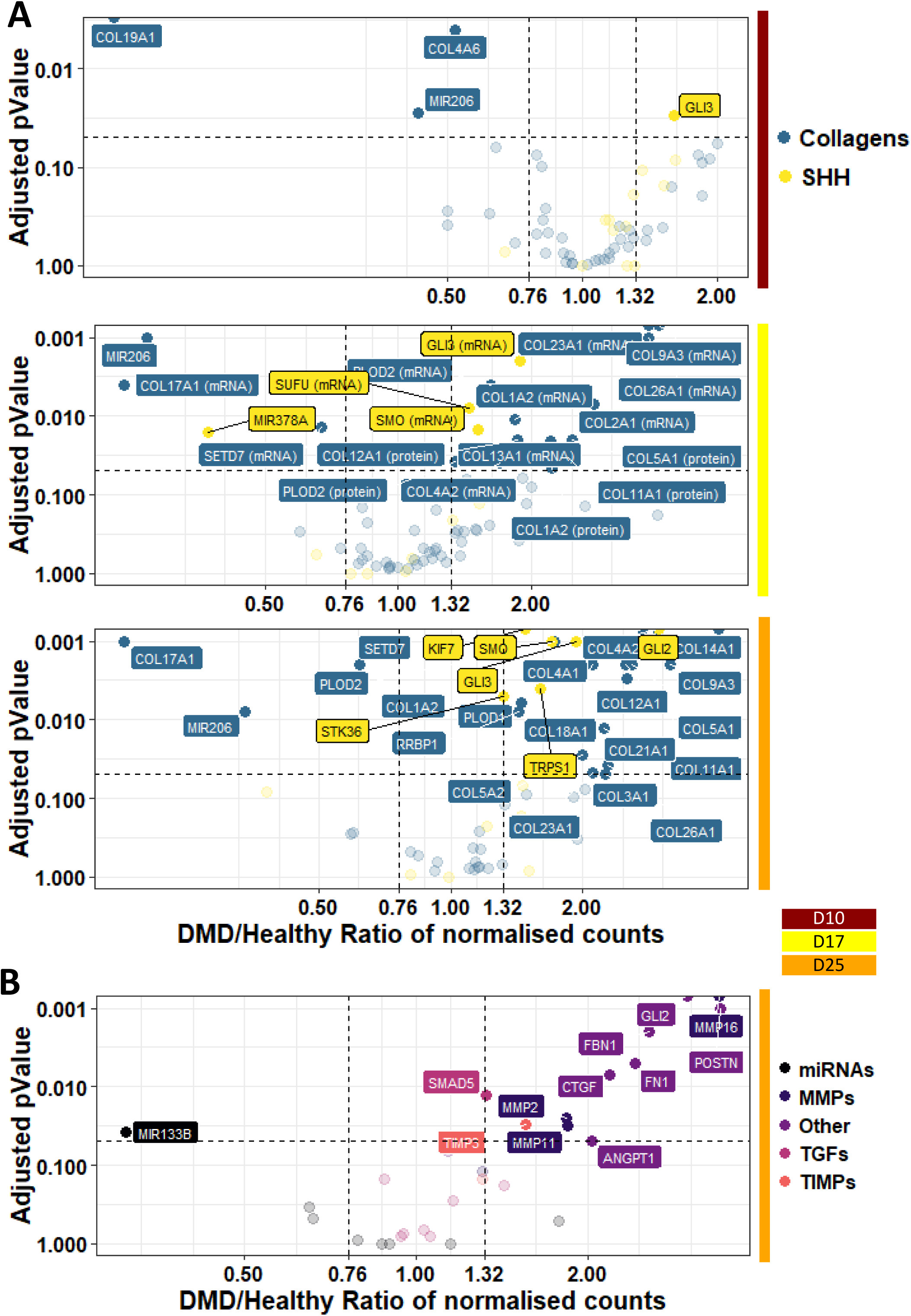
Illustration of the fibrosis phenotypes in DMD cells. Volcano plots of dysregulated mRNAs/miRNAs related to **A)** the SHH pathway and collagen metabolism at D10/17/25; and **B)** fibrosis at D25 – vertical grey dashed lines represent DMD/Healthy ratio thresholds at 0.76 or 1.32 - the horizontal grey dashed line represents the adjusted p-value threshold at 0.05. (D: day; MMP: matrix metallopeptidase; SHH: sonic hedgehog pathway; TIMP: tissue inhibitor of metallopeptidase; TGF: transforming growth factor).

At the myotube stage, a fibrosis-related gene set was clearly upregulated in DMD cells, as illustrated by the overexpression of *ANGPT1* (102), *CTGF* (103), collagens (*e.g. COL1A2* (104)), matrix metallopeptidases (*MMPs*) and tissue inhibitors of metallopeptidase (*TIMPs*) (105) (Figure 5B). Conversely, the myomiR *MIR133* that controls *CTGF* expression (106) was repressed (Table S2). Interestingly, gene members of the transforming growth factor (TGF)-β pathway, a well-known inducer of fibrosis (107), were not found dysregulated (Figure 5B, Table S2).

Altogether, these data argue for fibrosis as an intrinsic feature of DMD skeletal muscle cells, rather than a process solely driven by interstitial cell populations in the niche. Furthermore, this muscle-driven fibrosis seems independent of the TGF-β pathway, and could rather depend on the SHH pathway, together with an intrinsic upregulation of chondrocyte markers and collagens.

### Genes involved in mitochondrial metabolism are drastically dysregulated in DMD hiPSC-derived myotubes

As previously described (108) and illustrated on Figure S6A, genes involved in the energy metabolism of DMD hiPSC-derived myotubes were dysregulated at the creatine and carbohydrate levels, up to the respiration (Figure 6A-B, Figure S2C, Table S2). The creatine transporter was not impacted while mRNAs coding for enzymes of both creatine and creatine phosphate biosynthesis were underrepresented. Neither glucose nor glutamate transporter expression were impaired. However, genes involved in glutamine biosynthesis (followed by gluconeogenesis that feeds glycolysis from glutamine) as well as glycogenesis (followed by glycogenolysis that feeds glycolysis from glycogen) were all downregulated, together with genes coding for glycolysis itself. In contrast, genes coding for the pentose phosphate pathway (which is in parallel to glycolysis) were upregulated, especially the oxidative part. Gene expression for pyruvate decarboxylation and generation of acetyl-CoA to feed the tricarboxylic acid (TCA) cycle was also impaired. Finally, the genes involved in the TCA cycle itself (Figure 6A, Figure S2C) and the mitochondrial electron transport chain were downregulated Figure 6B, Figure S2C). This is particularly reinforced by lower levels of a member of the ATP synthase complex ATP5A1 at both mRNA and protein levels (Figure 6C-D). These mRNA and protein data were completed by the measurement of ATP levels, which were significantly decreased in DMD hiPSC-derived myotubes (Figure 6E). Moreover, transcripts encoded by the mitochondrial DNA and mitochondrial DNA itself were decreased in DMD hiPSC-derived myotubes at day 25 (Figure S6B-S6E).

**Figure 6.**
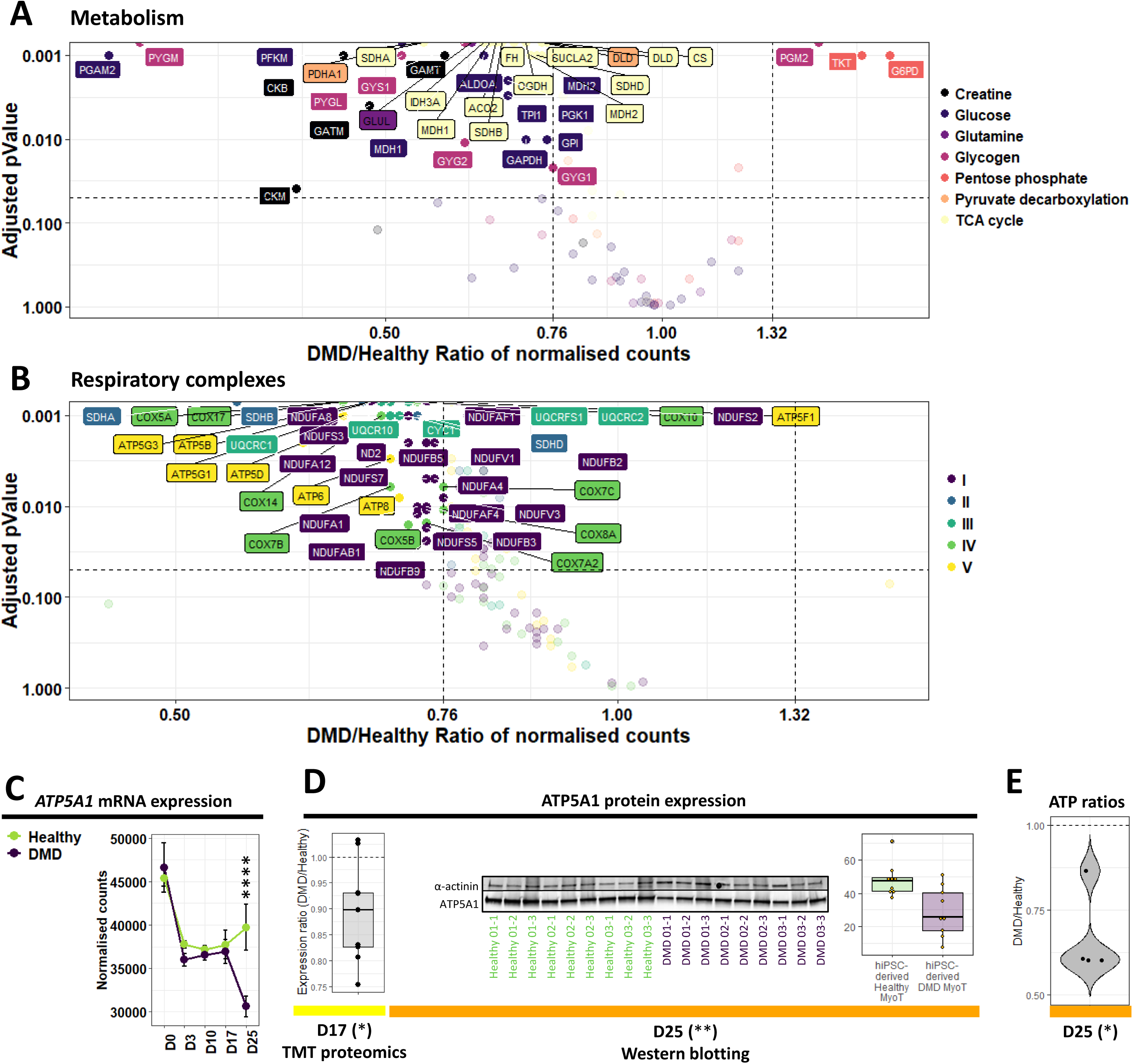
Illustration of the metabolic and mitochondrial phenotypes in DMD cells. Volcano plots of dysregulated mRNAs/miRNAs related to **A)** principal metabolic pathways; and **B)** the constitution of the five mitochondrial respiratory complexes in DMD hiPSC-derived MyoT – vertical grey dashed lines represent DMD/Healthy ratio thresholds at 0.76 or 1.32 - the horizontal grey dashed line represents the adjusted p-value threshold at 0.05. Quantification of ATP5A1 expression **C)** at the mRNA level during differentiation, and **D)** at the protein level at D17 (TMT proteomic data, left) and D25 (Western blot data, right). **E)** Measure of ATP levels in DMD cell lines, relative to Healthy controls. (*adjusted p-value ≤ 0.05, **adjusted p-value ≤ 0.01, ***adjusted p-value ≤ 0.001, ****adjusted p-value ≤ 0.0001). (D: day; hiPSC: human induced pluripotent stem cell, MyoT: myotube)

In the presented cell model, a significant downregulation of a mRNA set coding for mitochondrial proteins was primarily observed at day 10 with the downregulation of 11 % (12 mRNAs, DMD/Healthy expression ratio ≤ 0.76, adjusted p-value ≤ 0.05) of the mitochondrial outer membrane genes, and amplified during the differentiation of DMD cells (Figure 7A). Therefore, defects depicted at day 25 rooted before the expression of the skeletal muscle program at day 17. Among them, mRNA downregulation of *TSPO*, a channel-like molecule involved in the modulation of mitochondrial transition pore (109), occurred from day 10 to day 25. This downregulation was also observed at the protein level at day 17 (Figure 7B). Moreover, the protein import system was affected from day 17 at both mRNA and protein levels (Figure S6C-S6F). Simultaneously, mRNAs involved in mitochondrial genome transcription started to be downregulated, followed by genes involved in mitochondrial DNA replication at day 25 (Figure S6D-S6G). This progressive increase of dysregulations was also observed at the level of the entire mRNA set related to mitochondria (around 1,000 mRNAs) as illustrated by the volcano plots as well as the gene ontology enrichments (Figure 7C, Figure S2C).

**Figure 7.**
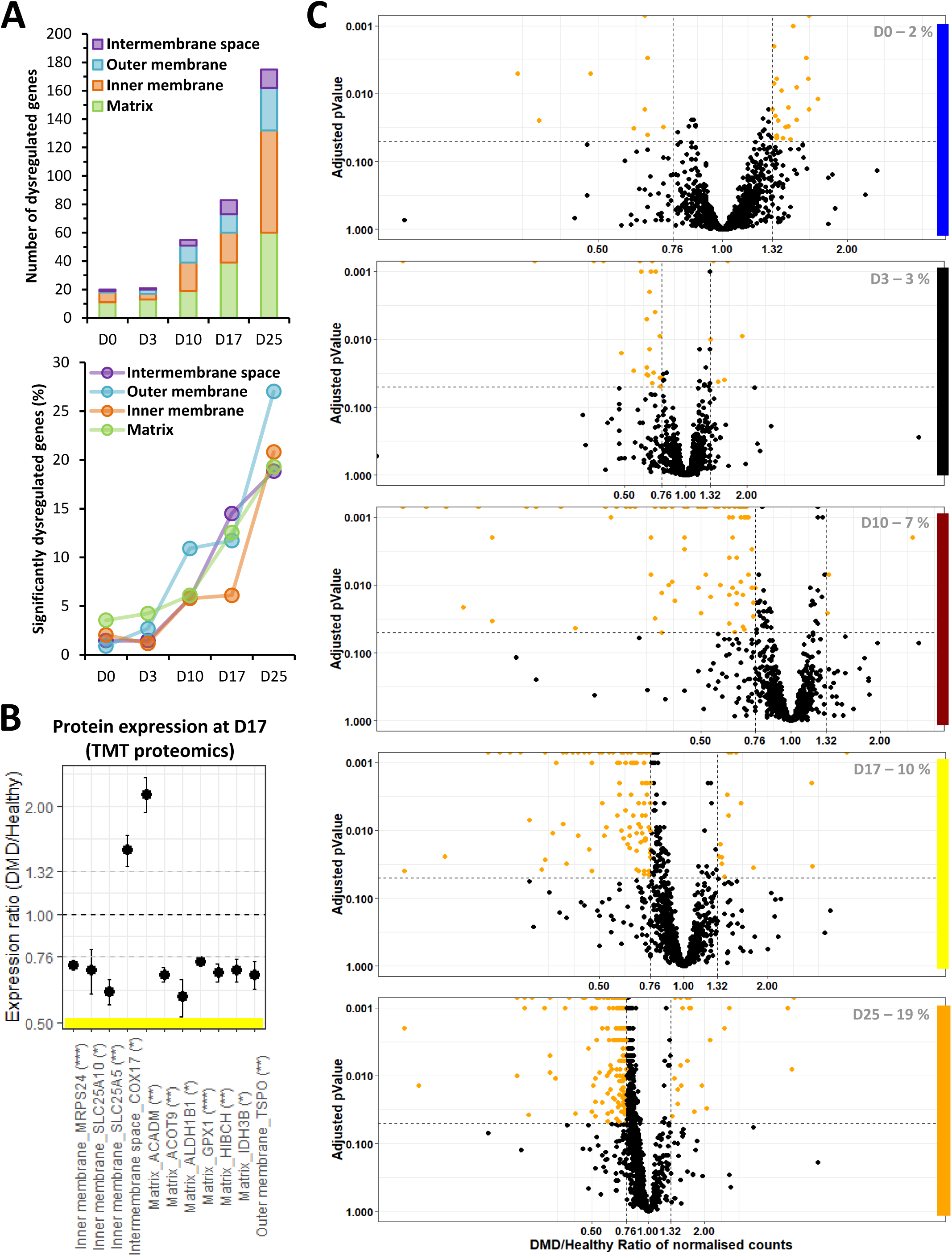
Mitochondrial dysregulations in DMD cells during differentiation. **A)** Absolute (top) and relative numbers (%, bottom) of dysregulated genes from the different mitochondrial compartments over the course of DMD hiPSC differentiation. **B)** Expression ratios of selected mitochondrial proteins. Statistical differences are indicated in brackets (*adjusted p-value ≤ 0.05, **adjusted p-value ≤ 0.01, ***adjusted p-value ≤ 0.001, ****adjusted p-value ≤ 0.0001). **C)** Volcano plots of mitochondria-related genes over the course of DMD hiPSC differentiation. Statistical differences are symbolised with orange dots – vertical grey dashed lines represent DMD/Healthy ratio thresholds at 0.76 or 1.32 - the horizontal grey dashed line represents the adjusted p-value threshold at 0.05 – The percentage of significantly dysregulated genes is indicated at the bottom right in grey. (D: day).

Our data highlight early impairments in genes coding for mitochondria that start at the somite stage and increase with the differentiation in an orderly manner. These elements complete the mitochondrial DMD phenotype described above at the myotube stage.

Altogether, our study demonstrates that DMD starts prior to the expression of well-described markers of muscle differentiation. It shows that hiPSC-based experimental models of DMD can help identify early disease manifestations and stratify multiple pathological features over the course of muscle development.

## DISCUSSION

Since the discovery of the *DMD* gene in 1987 (1), DMD cellular phenotypes were considered under the unique scope of a “mechanical hypothesis” in which dystrophin deficiency led to membrane leakage and ultimately muscle cell rupture. However, over the last 15-20 years, studies have brought unequivocal evidence that multiple additional factors are in play, such as calcium intracellular overloads (110,111), excessive oxidative stress (112,113), metabolic switches (114,115), as well as an overall tissue context where aberrant interactions between resident cells lead to inflammation and fibro-adipogenesis (116–118). This has progressively led to a complex picture involving interdependent homeostatic perturbations and to date, the identification of prevalent pathological features driving the initiation of DMD is hardly feasible.

The skeletal myogenesis modelled here by the differentiation of hiPSCs, without gene overexpression or cell sorting, homogeneously and robustly recapitulates key developmental steps – pluripotency, mesoderm, somite and skeletal muscle – without any trace of other lineages. Therefore, it is a suitable dynamic model for studying human skeletal muscle development in both healthy and DMD cells, offering the possibility to clarify the consequences of the absence of dystrophin at each step of the differentiation process, as well as to explore dystrophin functions and find earlier and more specific disease biomarkers.

As previously observed with pluripotent stem cells (119), hiPSC-derived myotubes at day 25 displayed an embryonic/foetal gene expression profile. However, a clear distinction must be made between the nature of the expressed isoforms – embryonic / foetal / postnatal – and the degree of differentiation. For instance, hiPSC-derived myotubes expressed multiple markers of terminally differentiated muscles at levels higher than those measured in tissue-derived myotubes. With the idea of exploring human DMD phenotypes during muscle development, we argued that generating embryonic/foetal myotubes from hiPSCs would not be a limitation.

In qualitative terms, DMD hiPSC-derived myotubes showed an overall morphology similar to healthy controls, with cell fusion and clear striation patterns, suggesting that the potential impact of dystrophin during *in vitro* differentiation is subtle and does not prevent myotube formation. However, our unbiased mRNA-seq analysis highlighted striking transcriptome dysregulations at day 25. This includes numerous genes which can be linked to previously described DMD phenotypes such as 1) DAPC dissociation (120); 2) rupture of calcium homeostasis (110); 3) myomiR downregulation (60,121); 4) sarcomere destabilisation (122–124); 5) mitochondrial and metabolism dysregulations (114,115); 6) NMJ fragmentation (125,126) and 7) fibrosis (118,127). It is interesting to note that these phenotypes are already detected at the transcriptional level in embryonic/foetal myotubes, while they usually appear postnatally in human patients and other animal models. In addition, most of them are often considered as consequences of degeneration-regeneration cycles typical of DMD muscles *in vivo* (123,128,129) which are absent in our *in vitro* model, indicating that a part of these defects are primarily due to the absence of dystrophin itself. In particular, our data suggest that fibrosis is an intrinsic feature of DMD skeletal muscle cells, and therefore, it does not absolutely require a specific tissue context or additional cell populations to be detected *in vitro*. Fibrosis is a major hallmark of DMD pathophysiology, and the regulation of this process has been largely investigated in the past (107,130). A long-debated question is the implication of the TGFβ signalling pathway (131). In this study, TGFβ signalling was inhibited up to day 17 by specific molecules contained in the cell culture media, and TGFβ-related genes were not upregulated at day 25, suggesting that the observed upregulation of fibrosis-related markers is TGFβ-independent.

Since several studies in human patients and animal models had described dystrophic phenotypes in DMD foetuses/infants (9–14), we investigated the precise timing of disease onset in our hiPSC-derived cells. First, the absence of dystrophin does not modify the capacity of cells derived from adult tissue biopsies to be reprogramed using the approach developed by Takeshi and Yamanaka (132). Both healthy and DMD cells retained pluripotency and the capacity to enter the mesoderm compartment at day 3. At that time, the embryonic dystrophin Dp412e is expressed and only marginal dysregulations are observed in DMD cells, *a priori* unrelated to cell fate choice as cells only express paraxial mesoderm markers at levels similar to healthy controls.

DMD dysregulations are greatly increased at day 10, when cells express somite markers. At that time, we noticed few significant dysregulations of cell lineage markers, which became more prevalent at day 17 and 25. This might be an indication that to some extent, cell fate is misguided in DMD cells, where skeletal muscle markers are underexpressed and replaced by markers of alternative lineages, such as chondrocytes. First visible at day 10, we identified the dysregulation of mitochondrial genes as one of the key processes happening in an orderly manner. Interestingly, early observations prior to the discovery of the *DMD* gene had hypothesised that DMD was a mitochondrial/metabolic disease based on protein quantifications and enzyme activities (114,133). Later, mitochondria was identified as a key organelle in DMD, responsible for metabolic perturbations but also calcium accumulation and generation of reactive oxygen species (110–113). In this study, numerous genes coding for proteins located in the outer mitochondrial membrane start to be downregulated from day 10 in DMD cells, such as the benzodiazepine receptor TSPO, a member of the controversial mitochondrial permeability transition pore (mPTP) (109). The mPTP is a multiprotein complex whose members are not all precisely identified, and several studies suggest that it might be involved in DMD pathophysiology (134,135). A chicken-and-egg question currently debated relates to the initiation of these homeostatic breakdowns, as positive feedbacks exist between mitochondria, oxidative stress and calcium homeostasis dysregulations (111,112). At the transcriptome level, dysregulations of genes controlling calcium homeostasis were detected after day 10, suggesting that mitochondrial impairment starts early and has predominant consequences in DMD, as hypothesised by Timpari *et al*. (108). Further experiments are needed to better evaluate the impact of mitochondrial dysregulations at the functional level.

Day 17 marks the entry into the skeletal muscle compartment with the expression of specific transcription factors, cell surface markers, myomiRs as well as the increase of skeletal muscle variant of dystrophin (*Dp427m*). It also marks the initiation of the skeletal muscle gene dysregulations observed at the myotube stage (*i.e.* downregulation of genes involved in DAPC and calcium homeostasis). For instance, the upregulation of fibrosis-related genes observed in DMD myotubes at day 25 is already visible at day 17, with the upregulation of the SHH pathway as well as collagen-related genes. In this study, it is seen as an early indicator of DMD physiopathology, confirming previous observations in DMD infants, both transcriptionally (4) and histologically (136,137).

Moreover, several myomiRs were found downregulated at days 17 and 25 and seem to play a central part in multiple DMD phenotypes. Beside their role in myogenesis (67,68), myomiRs can be involved in calcium homeostasis (138), metabolism and mitochondrial functions (139,140), and fibrosis (106,141). In particular, *MIR1-1* and *MIR206* are known to target key genes such as *CACNA1C* (138), *CTGF* (106), *RRBP1* (141), several regulators of the pentose phosphate pathway (139), and even transcripts encoded by the mitochondrial genome (140). Even though the functional consequences of the multiple gene and myomiR dysregulations highlighted in this study is virtually impossible to anticipate, we believe that myomiRs can be key players in DMD physiopathology.

Few studies argued that DMD starts before the expression of the muscular dystrophin protein (18,142). Our data suggests that *Dp427m* is actually expressed before muscle commitment but at a lower level. This fact might explain why disease phenotypes seem to be initiated at the somite stage. This early initiation could also be explained by the deficit in other dystrophin isoforms expressed before day 10, such as Dp412e at day 3 (15), as well as by the decrease or loss of other RNA products expressed from the *DMD* locus, such as the ubiquitous isoform Dp71-40 or long non-coding RNAs (143). The lack of knowledge around these additional products from the *DMD* locus contrasts with the extensive amount of data on the structure and function of the main muscular isoform Dp427m whose most studied role is to stabilise muscle cell membrane during contraction (144). *DMD* knockdown results at day 17 in a healthy cell line with partial mimicking of DMD phenotype could suggest a dynamic process in DMD: some dysregulations might not be reproduced by removing *DMD* after muscle commitment highlighting the fact that absence of DMD locus expression during development could have impacts before cells becoming muscles and, therefore, before Dp427m having its well-known role in muscles, as it is shown by our multi-omic study. The role of *Dp427m* in non-muscle cells could also be questioned. Other tissue specific isoforms have been described, *e.g.* in the retina (Dp260 (30)) and in the brain (Dp427c (145), Dp427p (29) and Dp140 (28)), some of which are also slightly expressed in skeletal muscles under certain circumstances (146), but their role remains mostly unknown. Interestingly, in our data, the expression of Dp260 follows the same pattern of expression as Dp427m. It has been shown that the expression of Dp260 in *mdx*/utrnK/K mice can rescue the *mdx* phenotype (147), indicating overlapping functions between Dp427m and Dp260. On the other hand, it is now well established that a third of DMD patients display cognitive deficiencies – which might be correlated with mutations affecting Dp140 (148) – attesting that dystrophin can be involved in other cell functions.

To date, the standard of care for DMD patients helps mitigate and delay some of the most severe symptoms but remains insufficient to have a curative effect. Despite decades of work with the *mdx* mouse model, only a few pharmacological candidate molecules have moved forward to clinical trials, with variable efficiency. As several gene therapy trials have been recently initiated with promising preliminary data, we believe that our human *in vitro* model system might be useful for the development of combination therapies. Recent studies have proved that the association of two different therapeutic approaches could have a synergistic effect on the overall treatment outcome, and can be used for instance to boost the effect of dystrophin re-expression by antisense oligonucleotides or gene therapy (8,149,150). Here, our extensive RNA-seq data could help identify relevant therapeutic targets for pharmacological intervention, such as CTGF – involved in fibrosis and found upregulated in DMD myotubes – which can be inhibited by monoclonal antibodies (151), or TSPO receptor – a receptor potentially member of the mPTP downregulated in DMD cells – targetable with benzodiazepines (152). In addition, our model might also be used as a platform to screen pharmacological compounds in an unbiased high-throughput manner. Indeed, skeletal muscle progenitor cells at day 17 can be robustly amplified, cryopreserved and plated in a 384-well plate format (data not shown). Thus, they could be an interesting tool to highlight pharmacological compounds to be used alone, or in combination with gene therapy.

To summarise, the directed differentiation of hiPSCs without gene overexpression or cell sorting homogeneously and robustly recapitulates key developmental steps of skeletal myogenesis and generates embryonic/foetal myotubes without any trace of other lineages. The absence of dystrophin does not compromise cell reprogramming, pluripotency or the entry into the mesoderm compartment. While a very low amount of the long muscular dystrophin isoform is expressed, a significant transcriptome dysregulation can be observed at the somite stage that implicates mitochondria prior to dysregulations of genes controlling calcium homeostasis. Despite their ability to enter the skeletal muscle lineage compartment and become myotubes, DMD cells exhibit an imbalance in cell fate choice as they express lower amounts of key muscle proteins and retain basal expression of marker genes from other lineages, leading to the well-characterised DMD phenotypes including muscle features and metabolism dysregulations as well as fibrosis. Altogether, these data argue for 1) a deficit and not a delay in DMD differentiation; 2) seeing DMD as a progressive developmental disease as well as a metabolic pathology whose onset is triggered before the entry into the skeletal muscle compartment; and 3) fibrosis as an intrinsic feature of DMD muscle cells. Future studies could explore the additional roles of *DMD* locus products and the impact of their loss during skeletal muscle development, as well as find earlier and more specific disease biomarkers and develop combination therapeutic strategies using high-throughput drug screening.

All the omics data from this study will be soon available online for exploration through a graphical interface. For additional information, please send an email to.

## MATERIALS AND METHODS

### Ethics, consent, and permissions

At the Cochin Hospital-Cochin Institute, the collection of primary cultures of myoblasts was established from patient muscle biopsies conducted as part of medical diagnostic procedure of neuromuscular disorders. For each patient included in this study, signed informed consent was obtained to collect and study biological resources, and establish primary cultures of fibroblasts and myoblasts at the Hospital Cell Bank-Cochin Assistance Publique—Hôpitaux de Paris (APHP). This collection of myoblasts was declared to legal and ethical authorities at the Ministry of Research (number of declaration, 701, n° of the modified declaration, 701–1) via the medical hosting institution, APHP, and to the “Commission Nationale de l’Informatique et des Libertés” (CNIL, number of declaration, 1154515).

### Cells

Human primary adult myoblasts from healthy individuals and DMD patients were provided by Celogos and Cochin Hospital-Cochin Institute (Table S3). In Celogos laboratory, cell preparation was done according to patent US2010/018873 A1.

### Cell culture

#### Human tissue-derived myoblasts

Primary myoblasts were maintained in a myoblast medium: DMEM/F-12, HEPES (31330–038, Thermo Fisher Scientific) supplemented with 10 % fetal bovine serum (FBS, Hyclone, Logan, UT), 10 ng/mL fibroblast growth factor 2 (FGF2, 100-18B, Peprotech), and 50 nM Dexamethasone (D4902, Sigma-Aldrich) on 0.1 % gelatin (G1393, Sigma-Aldrich) coated culture ware.

#### Human tissue-derived myotubes

Primary myoblasts were differentiated into myotubes. Cells were seeded at 600 cells/cm^2^ on 0.1 % gelatin coated cultureware in myoblast medium containing 1 mM Acid ascorbic 2P (A8960, Sigma-Aldrich).

#### Human induced pluripotent stem cells

Primary myoblasts were reprogrammed into hiPSCs following the protocol described in (15), using the Yamanaka’s factors POU5F1, SOX2 and KLF4 transduction by ecotropic or amphotropic vectors (Table S3). HiPSCs were adapted and maintained with mTeSR™1 culture medium (05850, Stemcell Technologies) on Corning^®^ Matrigel^®^ Basement Membrane Matrix, lactose dehydrogenase elevating virus (LDEV)-Free-coated cultureware (354234, Corning Incorporated). Cells were then seeded at 20,000 cells/cm^2^, passaged and thawed each time with 10 μM StemMACS™ Y27632.

#### Human iPSC-derived cell

*S*ix hiPSCs (3 healthy and 3 DMD) were differentiated three times toward skeletal muscle lineage using commercial media designed from Caron’s work (23) (Skeletal Muscle Induction medium SKM01, Myoblast Cell Culture Medium SKM02, Myotube Cell Culture Medium SKM03, AMSbio). This protocol is a 2D directed differentiation that uses 3 consecutive defined media (SKM01 from day 0 to 10, SKM02 from day 10 to 17 and SKM03 from day 17 to d25) and only one cell passage at day 10. Cells were seeded at 3,500 cells/cm^2^ at day 0 and day 10 on BioCoat™ Collagen I cultureware (356485, Corning Incorporated). Part of the cell culture was frozen at day 17 for further experiments such as DNA extraction. These cells were then thaw at 30,000 cells/cm^2^, and cultured in SKM02 for 3 days and SKM03 for 3 additional days to get myotubes.

### DNA and RNA experiments

#### RNA extraction and quality

RNA extraction was done in the six cell lines at 7 different time points: tissue-derived myoblast and tissue-derived myotube, as well as during hiPSC differentiation at day 0, 3, 10, 17 and 25 (hiPSC-derived myotube) using the miRNeasy Mini kit (217004, QIAgen) on the QIAcube instrument. RNAs coming from the part A of the extraction protocol was used for mRNA-seq and RT-qPCR. RNAs coming from the part B of the extraction protocol was used for miRseq. PartA RNA was quantified on Nanodrop spectrophotometer (ND-1000, Thermo Fisher Scientific) and purity/quality (RIN ≥ 7) was assessed on the 2200 TapeStation using the Agilent RNA ScreenTape (5067-5576 / 5067-5577 / 5067-5578, Agilent). PartB RNA was quantified and purity/quality was assessed on the 2100 Agilent BioAnlayzer using the Agilent small RNA kit (5067-1548, Agilent).

#### Reverse transcription

500 ng of total RNA were reverse transcribed with random primers (48190–011, Thermo Fisher Scientific), oligo(dT) (SO131, Thermo Fisher Scientific), and deoxynucleotide (dNTP, 10297–018, Thermo Fisher Scientific) using Superscript^®^ III reverse transcriptase (18080–044, Thermo Fisher Scientific). Thermocycling conditions were 10 min, 25 °C; 60 min, 55 °C; and 15 min, 75 °C.

#### qPCR

We amplified cDNA/total DNA using primers (Thermo Fisher Scientific) listed in Table S6. They were designed using Primer blast (http://www.ncbi.nlm.nih.gov/tools/primer-blast). The amplification efficiency of each primer set was preliminarily determined by running a standard curve. Detection was performed using a QuantStudio™ 12K Flex Real-Time PCR System (Thermo Fisher Scientific). Reactions were carried out in a 384-well plate, with 10 μL containing 2.5 µL of 1/10 cDNA or 6.25 ng/uL total DNA, 0.2 µL of mixed forward and reverse primers at 10 µM each, and 5 µL of 2X Luminaris Color HiGreen qPCR Master Mix Low Rox (K0973, Thermo Fisher Scientific). Thermocycling conditions were 50 °C during 2 min, 95 °C during 10 min, followed by 45 cycles including 15 sec at 95 °C, 1 min at 60 °C plus a dissociation stage. All samples were measured in triplicate. Experiments were normalised using UBC as reference gene and relative quantification was done with the ΔΔCt method.

#### mRNA-seq

Libraries are prepared with TruSeq Stranded mRNA kit protocol according supplier recommendations. Briefly, the key stages of this protocol are successively, the purification of PolyA containing mRNA molecules using poly-T oligo attached magnetic beads from 1µg total RNA, a fragmentation using divalent cations under elevated temperature to obtain approximately 300bp pieces, double strand cDNA synthesis and finally Illumina adapter ligation and cDNA library amplification by PCR for sequencing. Sequencing is then carried out on paired-end 100 b/75 b of Illumina HiSeq 4000.

An RNA-seq analysis workflow was designed using snakemake 3.5.4 (153) for read quality estimation, mapping and differential expression analysis. Quality estimation was obtained with FastQC 0.11.5 (https://www.bioinformatics.babraham.ac.uk/projects/fastqc/). Mapping to the human genome assembly Ensembl GRCh37.87 (43,695 transcripts) was performed with STAR 2.5.0a (154). According to STAR manual and for more sensitive novel junction discovery, the junctions detected in a first round of mapping were used in a second mapping round. Read strandness was confirmed using RSeQC (155). Analysis results were summarised using MultiQC 1.0 (156). Normalised counts (median ratio normalisation, MRN) and differential expression analysis was performed with DESeq2 1.16.1 (157), considering pairwise comparisons with all developmental stages and comparing DMD versus healthy cells within developmental stages. BiomaRt 2.30.0 (158) was used to fetch gene annotations from Ensembl. Transcripts with |log2FoldChange| ≥ 0.4 (equivalent of DMD/healthy ratio ≤ 0.76 or ≥ 1.32) and adjusted p-value ≤ 0.05 were considered differentially expressed. RNA-seq data have been deposited in the ArrayExpress database (159) at EMBL-EBI under accession number E-MTAB-8321 (https://www.ebi.ac.uk/arrayexpress/experiments/E-MTAB-8321).

#### miRNA-seq

10 ng of miRNA was reverse transcribed using the Ion Total RNA-seq kit v2 (4475936, Thermofisher Scientific) following the protocol of the manufacturer for small RNA libraries. The cDNA libraries were amplified and barcoded using Ion Total RNA-seq kit v2 and Ion Xpress RNA-seq Barcode Adapters 1-16 Kit (Thermofisher Scientific). The amplicons were quantified using Agilent High Sensitivity DNA kit before the samples were pooled in sets of fifteen. Emulsion PCR and enrichment was performed on the Ion OT2 system Instrument using the Ion PI Hi-Q OT2 200 kit (A26434, Thermofisher Scientific). Samples were loaded on an Ion PI v3 Chip and sequenced on the Ion Proton System using Ion PI Hi-Q sequencing 200 kit chemistry (200 bp read length; A26433, Thermofisher Scientific). Sequencing reads were trimmed with Prinseq (160) (v0.20.4) (--trim-right 20) and filtered by average quality score (--trim-qual 20). Reads with a size less than 15 bp have been removed and reads with a size greater than 100 bp have been trimmed with Cutadapt (v1.16)(161). Mapping to the human genome assembly Ensembl GRCh37.87 (3111 transcripts) was performed with STAR 2.5.3a (154). Normalised counts (median ratio normalisation, MRN) and differential expression analysis was performed with DESeq2 1.16.1 (157), considering pairwise comparisons with all developmental stages and comparing DMD versus healthy cells within developmental stages. Transcripts with |log2FoldChange| ≥ 0.4 (equivalent of DMD/healthy ratio ≤ 0.76 or ≥ 1.32) and p-value ≤ 0.05 were considered differentially expressed. The use of p-value instead of adjusted p-value is justified by biological meaning(162) (i.e. well-known regulated / dysregulated miRNAs had a p-value ≤ 0.05 but not an adjusted p-value ≤ 0.05). miRNA-seq data have been deposited in the ArrayExpress database (159) at EMBL-EBI under accession number E-MTAB-8293 (https://www.ebi.ac.uk/arrayexpress/experiments/E-MTAB-8293).

#### High-throughput data analyses

Graphs were realised using RStudio. Viridis 0.5.1 library (163) was used for the colour palette easier to read with colour blindness and print well in grey scale. For unsupervised analyses, normalised counts were standardised with scale function (center = TRUE, scale = TRUE) and plotted with corrplot function from corrplot 0.84 library (164). Spearman correlation was done with the cor function (method = “spearman”, use = “pairwise.complete.obs”) on standardised data. Hierarchical clustering and heatmap were performed with gplots 3.0.3 library (165) heatmap.2 function on standardised data. Gene enrichment data were retrieved from DAVID database using RDAVIDWebService 1.24.0 library (166) on supervised list of mRNAs (mRNA-seq data: adjusted p-value ≤ 0.01, normalised counts ≥ 5 in at least one sample, ratio ≤ 0.5 or ≥ 2 for myogenesis (Figure S2B) and ratio ≤ 0.76 or ≥ 1.32 for DMD phenotype (Figure S2C); enrichment data: Benjamini value ≤ 0.05, enrichment ≥ 1.5). Only Gene Ontology terms were processed. Spearman correlations for the comparison transcriptomics vs proteomics at day 17 and for comparisons with published omics datasets were performed using two-tailed nonparametric Spearman correlation on GraphPad Prism software.

#### Exon skipping

1,000,000 healthy M180 cells were transfected after 17 days of culture by electroporation with a phosphorodiamidate morpholino oligo (PMO) targeting exon 7 of the *DMD* gene at 10 or 100 µM, or a PMO Control at 100 µM in 100 µL solution from the P3 Primary Cell 4D-Nucleofector^®^ X Kit L (V4XP-3024, Lonza) using the CB150 program on the 4D-Nucleofector^™^ System (Lonza). Cells were seeded at a density of 100,000 cells/cm^2^. RNA extraction was carried on transfected cells 24 h, 48 h and 72 h later followed by RT as described above. PCR was done on1 uL of cDNA using 10 µM of forward and reverse primers (Fw 5’-AAGATTCTCCTGAGCTGGGTC −3’ and Rv 5’-AGTCACTTTAGGTGGCCTTGG −3’, Life technologies) and 1 U Taq DNA polymerase (10342, Life technologies) as described by the manufacturer’s instructions, for a final reaction volume of 25 µL. PCR reaction started by a step at 94°C for 3 min, followed by 27 cycles at 94°C for 45 s, 55°C for 45 s and 72°C for 45 s, and a final step at 72°C for 5 min. Exon skipping was analyzed using the DNA 1000 kit (5067, Agilent) on the Agilent 2100 Bioanalyzer. Full length PCR product was 372 bp and exon skipped length PCR product was 253 bp. Results were computed by the Agilent 2100 Bioanalyzer software v3.81. Spearman correlations were performed using two-tailed nonparametric Spearman correlation on GraphPad Prism software.

### Protein experiments

#### Immunolabelling

Cells (healthy hiPSC 1/ DMD hiPSC 2, Table S5) at day 17 of culture were thawed and seeded at 10,000 cells/cm^2^ in SKM02 medium in Falcon^®^ 96-well microplate (353219, Corning) coated with 0.1% gelatin (G1393, Sigma-Aldrich) and 2.4 μg/mL laminin (23017015, Thermofischer Scientific) in PBS 1X (D8537, Sigma-Aldrich). After 4 days, cells were switched to DMEM/F-12, HEPES (31330038, Thermofischer Scientific) with 2% Horse serum (H1270, Sigma-Aldrich). Before staining, after removing the culture medium, cells were fixed 15 min at 4°C with PFA 4% (15710, Euromedex) after 7 days of culture. A first quick Phosphate buffered saline (PBS) 1X tablets (P4417, Sigma-Aldrich) wash was done, followed by another lasting 10 min. Then, a solution with PBS 1X, Triton™ X-100 0.25% (T8787, Sigma-Aldrich) and Bovine serum albumin 2.5% (BSA, A9418, Sigma-Aldrich) was added and incubated 30 min at room temperature. Primary antibody was finally added, diluted in the same buffer (α-actinin 1/500, A7811, Sigma-Aldrich), overnight at 4°C. The next day, two quick PBS 1X washes were followed by a third incubated 10 min at room temperature. An incubation was done 45 min at room temperature with a mix of 4′,6-Diamidine-2′-phenylindole dihydrochloride (DAPI, 1µg/mL, 10236276001, Sigma-Aldrich) and the secondary antibody Donkey anti-Mouse Alexa Fluor 555 in PBS 1X, (1/1000, A-31570, Thermofischer Scientific). Finally, two quick PBS 1X washes were followed by a third incubated 10 min at room temperature. The stained cells were kept in PBS 1X at 4°C before imaging with a Zeiss LSM880 Airyscan confocal and Zen software (Black edition).

#### Western blotting

For tissue-derived myotubes, after three rinses with cold PBS 1X (w/o Ca2+ and Mg2+, D8537, Sigma-Aldrich), protein extracts were isolated from cultured cells by scraping (010154, Dutscher) with an extraction protein buffer (NaCl 150 mM, Tris 50 mM, EDTA 10 mM (AM9260G, ThermoFisher Scientific), Triton 1X, 1/100 Protease Inhibitor Cocktail (P8340, Sigma-Aldrich), PhosSTOP tablet (04906845001, Roche Diagnostics)). For hiPSC-derived myotubes, cell pellets were rinsed once with cold PBS 1X, spun 5 min at 300 g and resuspended in the same extraction protein buffer. Protein Extracts were centrifuged at 4°C 10 min at 16,000 g and supernatants were kept at −80 °C. Quantitation of total protein was done with Pierce BCA protein assay kit (23225, ThermoFischer Scientific). Before gel loading, protein extracts were mixed with 9µL of loading buffer (Urea 4M, SDS 3.8%, Glycerol 20%, Tris 75mM pH 6.8, 5% β-mercaptoethanol, 0.1mg/mL Bromophenol blue) and completed to 28µL (for one well) with extraction protein buffer, then heated once 5 min at 95 °C. Western blots were performed either with Criterion ™ XT Tris-Acetate Precast Gels 3–8 % (3450130, Bio-Rad, Hercules, CA), XT Tricine running buffer (161–0790, Bio-Rad) and ran at room temperature for 1 hour and 15 min at 150 V for RYR1 (1/1000, MA3-925, ThermoFisher Scientific), MF20 (1/500, DSHB, concentrate), Manex50 (1/30, DSHB), α-sarcoglycane (1/150, A-SARC-L-CE, Leica biosystems), γ-sarcoglycane (1/150, G-SARC-CE, Leica biosystems), or with 4–15% Criterion™ TGX™ Precast Midi Protein Gel (5671084, Bio-Rad), 10x Tris/Glycine/SDS Running Buffer (1610772), and ran at room temperature for 1 hour at 200 V for CaV1.1 (1/1000, MA3-920, ThermoFisher Scientific), ATP5A (1/1,000, ab14748, ABCAM), Semaphorin 6A (1/55, AF1146, R&D systems) and GLI3 (1/200, AF3690, R&D systems). Gels were rinsed once in water and blotted either with “high molecular weight” or “mixed molecular weight” program of TransBlot^®^ Turbo™ transfer system (Bio-Rad) using Trans-Blot^®^Turbo™ Midi Nitrocellulose Transfer Packs (170–4159, Bio-Rad). Blots were then processed with the SNAP i.d.^®^ 2.0 Protein Detection System following the manufacturer’s protocol, with Odyssey^®^ Blocking Buffer (927-40003, LI-COR) for blocking and with 0,2% Tween^®^ 20 added for antibody dilutions (28829.296, VWR), washes were done with phosphate-buffered saline tween (PBST) buffer (PBS 1X tablets, P4417, Sigma-Aldrich; 0.1 % Tween^®^ 20). Every primary antibody was pooled with either α-actinin (1/12,500, sc-17829, Santa Cruz or 1/7000, A7811, Sigma-Aldrich) or α-tubulin (1/6666, Ab7291, Abcam). For secondary antibodies, either IRDye 800CW donkey anti-mouse and/or IRDye^®^ 680RD donkey anti-goat were used (1/5000-1/10000, 926-32212, 926-68074, LI-COR). After completion of SNAP i.d.^®^ general protocol, with the membrane still in the blot holder, two PBS 1X washes were finally done before band visualisations with Odyssey^®^ CLx Imaging System and quantification with Image Studio Lite software (Version 5.2). Statistical analysis was performed using unpaired t test on GraphPad Prism software.

#### TMT Isobaric quantitative proteomics

##### Samples Preparation

Cells at day 17 were collected and resuspended in 90% FBS (Hyclone), 10% DMSO (A3672.0050, VWR), cooled down until −90°C with the CryoMed™ device (ThermoFisher Scientific), before storage in liquid nitrogen. Cells were then thawed and washed 5 times with cold PBS and air was replaced by Argon to thoroughly dry the pellet that was flash frozen in liquid nitrogen. 5-10 times the approximate cell pellet volume of 0.5 M triethyl ammonium bicarbonate (TEAB) with 0.05% SDS was added to the cell pellet for protein extraction. Cell pellet was re-suspended and triturated by passing through a 23-gauge needle and 1ml syringe for 30 times. Samples were then sonicated on ice at amplitude of 20% for 30 × 2 sec bursts and centrifuged at 16000g for 10 min at 4°C. Supernatant was transferred to a fresh Eppendorf tube. Protein was quantified by nanodrop. 100-150µg of protein was aliquoted for each individual sample and 2µl TCEP (50mM tris-2-carboxymethyl phosphine) was added for every 20µl of protein used for reducing the samples. After 1 hr incubation at 60°C, 1µl MMTS (200mM methylmethane thiosulphonate) was added for every 20µl of protein used for alkylating/’blocking’ the samples. Finally, after a 10 min incubation at RT, samples were trypsinised by addition of 6-7.5µl of 500ng/µl trypsin. The ration between enzyme: substrate was 1:40. Samples were incubated overnight at 37°C in the dark.

##### TMT labelling

When TMT reagents reached room temperature, 50µl of isopropanol/[acetonitrile] was added to each TMT 11-plex reagent and was incubated at RT for 2 hrs, in the dark. 8 µl of 5% hydroxylamine was added to neutralise the reaction. Each sample was separately lyophilised at 45°C. Samples have been stored at −20°C or used immediately.

##### Offline C4 High Performance Liquid Chromatography (HPLC)

All 8 samples were pooled together in 60µl of 97% mobile phase A (99.92% % H2O, 0.08% NH_4_OH) and 3% mobile phase B (99.92% % Acetonitrile, 0.02% NH_4_OH) by serially reconstituting each sample. Extra 40µl of mobile phase was added to sample 1, after sample has been well vortexed, all the contents of sample 1 tube were transferred to the tube with the sample 2 (and serially repeated until all samples were pooled). Final volume of samples needed to be 100µl. After sample was centrifuged at 13000g for 10 min, supernatant was collected with an HPLC injection syringe. 100µl was injected onto the sample loop. Fractions were collected in a peak dependent manner. Finally, fractions were lyophilised at 45°C and stored at −20°C until required. The used column was a Kromasil C4 column 100Å pore size, 3.5µm particle size, 2.1mm inner diameter and 150mm length. The gradient for C4 separation was (RT in min - %B): 0-3; 10-3; 11-5; 16-5; 65-20; 100-30; 15-80; 120-80; 125-3.

##### Solid Phase Extraction Cleaning of peptides fractions

A GracePureTMT SPE C18-Aq cartridge was used for pre-cleaning of samples (Support: Silica, % Carbon: 12.5%, With endcapping, Surface area: 518m^2^/g, Particle size: 50μm, Pore size: 60Å, Water-wettable). Samples were reconstituted using in total 400μl of 1% ACN, 0.01% FA. Cartridge was washed with 600μl of ACN. ACN was then completely flushed out of the column at dropwise speed. This activated the ligands. Then 1% ACN, 0.01% FA (600μl) was flushed through the cartridge to equilibrate the sorbent. 400μl of the sample was loaded in the cartridge. It was then very slowly flushed through the cartridge and recovered into a fresh tube. This process was repeated 3 times. 2 volumes of 250μl of 1%ACN, 0.01%FA were used to clean and de-salt the sample. It was flushed through very slowly. 2 volumes (250μl each) were used per step (2% ACN, 10% ACN, 30% ACN, 50% ACN, 70% ACN). This cycle was repeated twice. Each particular concentration was pooled in one tube. Samples were dried to dryness in a Speedvac at RT overnight and stored at −20°C. Like previously, samples were pooled with 100µl of 97% mobile phase A (99.92% % H2O, 0.08% NH_4_OH) and 3% mobile phase B (99.92% % Acetonitrile, 0.02% NH_4_OH) and injected onto the sample loop. Fractions were collected in a peak dependent manner. The gradient for SPE cleaned peptides C4 separation (RT in min - %B): 0-2; 10-2; 20-5; 25-5; 35-20; 55-35; 60-35; 70-80; 75-80; 80-3.

##### Online C18 High Precision Liquid Chromatography (HPLC)

30µl of loading phase (2% acetonitrile, 1.0% formic acid) was added to each fraction-containing Eppendorf tube. Samples were vortexed and centrifuged. Blanks (30µl mobile phase) were added into well A1 to A12. 30µl of sample 1 was pipetted into well B1, sample 2 in well B2 and so on. An orthogonal 2D-LC-MS/MS analysis was performed with the Dionex Ultimate 3000 UHPLC system coupled with the ultra-high-resolution nano ESI LTQ-Velos Pro Orbitrap Elite mass spectrometer (Thermo Scientific).

##### Data analysis

HCD and CID tandem mass spectra were collected and submitted to Sequest search engine implemented on the Proteome Discoverer software version 1.4 for peptide and protein identifications. All spectra were searched against the UniProtKB SwissProt. The level of confidence for peptide identifications was estimated using the Percolator node with decoy database searching. False discovery rate (FDR) was set to 0.05, and validation was based on the q-Value. Protein ratios were normalised to protein median and peptides with missing TMT values were rejected from protein quantification. Phosphorylation localisation probability was estimated with the phosphoRS node. Protein ratios were transformed to log_2_ ratios and significant changes were determined by one sample T-test. To reduce the impact of possible false positive identifications, more parameters were set: 1) only proteins with more than two quantified unique peptides. 2) DMD/Healthy ratio ≥ 1.32 or ≤ 0.76 and 3) only FDR corrected p-value ≤ 0.05 were retained for bioinformatics analysis. The list of proteins quantified in the 6 samples is in Table S3. Proteomic data have been deposited in the PRIDE Archive database (167) at EMBL-EBI under accession number PXD015355 (https://www.ebi.ac.uk/pride/archive/projects/PXD015355).

#### ATP experiments

Two healthy (M180 and M398) and two DMD (M202 and M418) cell lines after 17 days of culture were seeded in 384-well plates at a density of 30,000 cells/cm^2^. Living cells were staining with HOECHST at a concentration of 1/300 six days later for cell quantification (nuclei per well were counted using the CX7 imaging system, ThermoFisher). ATP measure was done using the CellTiter-Glo™ Luminescent Cell Viability Assay Kit (Promega) following the manufacturer’s protocol and normalised by the cell quantification. Statistical analysis was performed using one-sample t test on GraphPad Prism software (each healthy cell line was compared to each DMD cell line).

## ACKNOWLEDGEMENTS

The RNA-Sequencing libraries were processed and sequenced by Integragen (Evry, France). We gratefully acknowledge support from the PSMN (Pôle Scientifique de Modélisation Numérique) of the ENS de Lyon for the computing resources. We thank Dr Nacira Tabti, Dr Elisabeth Le Rumeur, Dr Nathalie Deburgrave and Dr Malgorzata Rak for providing us with specific reagents and antibodies. We thank Dr Linda Popplewell for designing and validating the PMO7 sequence. We thank Dr David Israeli for his feedback on the manuscript and overall discussion on our project.

## FIGURE LEGENDS

**Figure S1 – DMD variant expression over the course of hiPSC differentiation. A)** Bright field microscope pictures at the 7 differentiation points giving rise to hiPSC-derived and tissue-derived MyoT. Possible cryopreservation time points are indicated by snowflakes. **B)** RT-qPCR relative quantification of *DMD* variants expression during differentiation of hiPSCs (D0) into MyoT (D25) with the related cycle threshold (CT) values (Ct: cycle threshold; D: day; hiPSC: human induced pluripotent stem cell; MyoB: myoblast; MyoT: myotube).

**Figure S2 – Gene ontology enrichments over the course of healthy and DMD hiPSC differentiation A)** Proportions of significantly regulated mRNAs (adjusted p-value ≤ 0.01) between successive differentiation time points during the differentiation of healthy hiPSCs. Gene ontology enrichments on **B)** significantly regulated mRNAs between successive differentiation time points in healthy cells (number of genes in brackets) and **C)** significantly dysregulated mRNAs at each differentiation time points in DMD cells. The number of genes involved in these significant enrichments is indicated in brackets next to each GO term. In green, GO terms related to downregulated genes and in yellow, GO terms related to upregulated genes (BP: biological process; CC: cellular component; D: day; hiPSC: human induced pluripotent stem cell; MyoB: myoblast; MyoT: myotube).

**Figure S3 – Comparison of healthy and DMD cells at D10 and D17, protein analyses.** Western blots and quantifications of **A)** SEMA6A at D10, **B)** GLI3 at D10 and **C)** GLI3 at D17. Omics comparison of mRNA and protein data at day 17: **D)** Venn diagram of the number of genes with |log2FoldChange| ≥ 0.4 and adjusted pvalue ≤ 0.05 in either transcriptomic or proteomic data, **E)** their associated Spearman correlation coefficient in brown, as well as **D)** their correlation graph with the number of genes with |log2FoldChange| ≥ 0.4 in both sets are indicated (genes with p-value ≥ 0.05 only in transcriptomics are in blue, only in proteomics in purple and in both in orange). (*p-value ≤ 0.05, **p-value ≤ 0.01, ***p-value ≤ 0.001, ****p-value ≤ 0.0001; D: day; GLI3FL: GLI3 full length; GLI3R: GLI3 repressor).

**Figure S4 – *DMD* knockdown at D17 in healthy cells. A)** qPCR quantification of *DMD* expression related to exon skipping efficiency (%); **B)** Western blot quantification of dystrophin; and **C)** qPCR quantification of selected genes following exon skipping (boxplots of expression ratio of exon 7 skipped/unskipped conditions when the exon skipping efficiency was above 70% at the top, and Spearman correlation between all skipped and unskipped conditions at the bottom; ****p-value < 0.0001, ns: not significant).

**Figure S5 – Comparison of hiPSC-derived and tissue-derived MyoT for the expression of cell cycle genes and myogenic regulators.** Hierarchical clustering and heatmap of **A)** selected cell cycle transcripts and miRNAs, and **B)** DLK1, IGF2 and selected myosin transcripts in hiPSCs (D0), hiPSC- and tissue-derived MyoT. **C)** Dotplot of DMD/healthy expression ratio of muscle transcription factors. Significant statistical differences are shown in brackets (*adjusted p-value ≤ 0.05, **adjusted p-value ≤ 0.01, ***adjusted p-value ≤ 0.001, ****adjusted p-value ≤ 0.0001). (hiPSC: human induced pluripotent stem cell; MyoT: myotube).

**Figure S6 – Dysregulations of metabolic pathways and mitochondrial genes during differentiation of DMD hiPSCs. A)** Scheme of metabolism dysregulations at day 25. Dotplots of **B)** mitochondrial transcripts, **C)** transcripts coding mitochondrial protein import, and **D)** transcripts coding mitochondrial transcription/replication; **E)** Mitochondrial DNA quantification by qPCR at D25. Dotplots of mitochondrial proteins expressed at D17 involved in **F)** protein import, **G)** mitochondrial transcription/replication. Statistics are in brackets (*adjusted p-value ≤ 0.05, **adjusted p-value ≤ 0.01, ***adjusted p-value ≤ 0.001, ****adjusted p-value ≤ 0.0001; D: day).

