## Supplemental Figures for "Myogenesis modelled by human pluripotent stem cells uncovers Duchenne muscular dystrophy phenotypes prior to skeletal muscle commitment"

### Figure S1

**A**

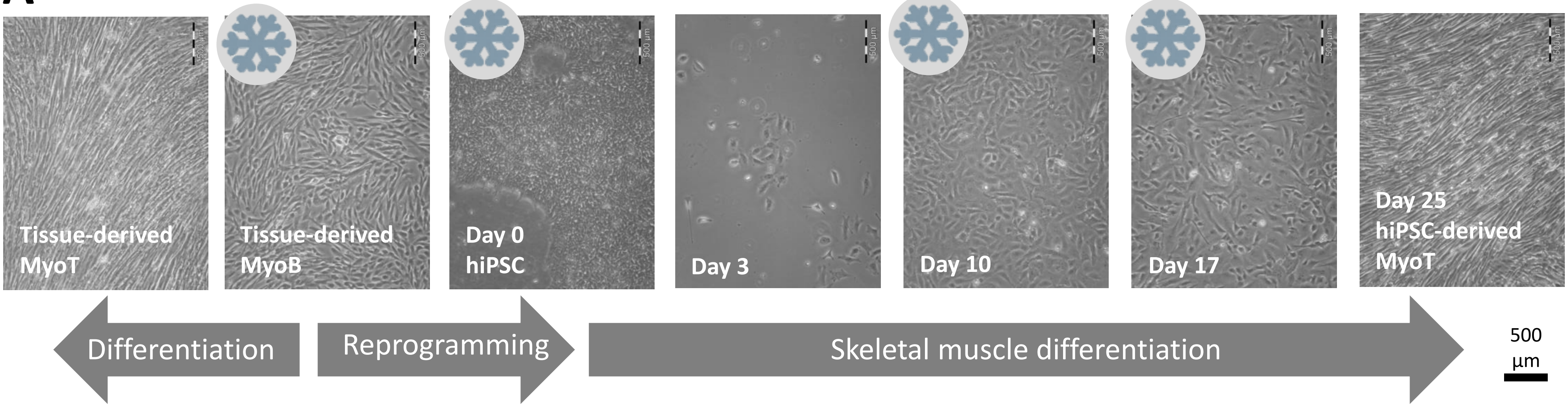


**B**

| Target Name | Cell state | Phenotype | Ct mean | Ct SEM |
|---|---|---|---|---|
| ***UBC*** (reference gene) | D0 | Healthy | 23.7 | 0.1 |
| | | DMD | 24.5 | 0.3 |
| | D3 | Healthy | 24.9 | 0.3 |
| | | DMD | 24.8 | 0.3 |
| | D10 | Healthy | 24.9 | 0.2 |
| | | DMD | 25.5 | 0.4 |
| | D17 | Healthy | 25.7 | 0.6 |
| | | DMD | 26.0 | 0.7 |
| | D25 | Healthy | 25.8 | 0.4 |
| | | DMD | 27.1 | 1.4 |
| ***Dp71-40*** | D0 | Healthy | 25.2 | 0.3 |
| | | DMD | 26.1 | 0.0 |
| | D3 | Healthy | 25.2 | 0.2 |
| | | DMD | 26.4 | 0.3 |
| | D10 | Healthy | 24.5 | 0.7 |
| | | DMD | 24.5 | 0.2 |
| | D17 | Healthy | 27.2 | 0.5 |
| | | DMD | 26.5 | 0.4 |
| | D25 | Healthy | 26.6 | 0.8 |
| | | DMD | 25.9 | 1.9 |
| ***Dp116*** | D3 | Healthy | 35.9 | 0.3 |
| | D10 | Healthy | 33.6 | 2.3 |
| | | DMD | 35.0 | 0.5 |
| | D17 | Healthy | 37.9 | 2.6 |
| | | DMD | 35.6 | 0.4 |
| | D25 | Healthy | 34.2 | 1.2 |
| | | DMD | 35.3 | 2.5 |
| ***Dp140*** | D0 | Healthy | 32.1 | 1.6 |
| | | DMD | 33.8 | 0.7 |
| | D3 | Healthy | 32.7 | 0.7 |
| | | DMD | 36.5 | 0.2 |
| | D10 | Healthy | 32.7 | 2.8 |
| | | DMD | 34.4 | 0.7 |
| | D25 | Healthy | 36.5 | 0.2 |
| ***Dp260*** | D0 | Healthy | 31.2 | 0.5 |
| | | DMD | 31.6 | 1.0 |
| | D3 | Healthy | 29.3 | 1.4 |
| | | DMD | 30.0 | 2.3 |
| | D10 | Healthy | 30.0 | 1.0 |
| | | DMD | 30.3 | 0.6 |
| | D17 | Healthy | 25.3 | 1.2 |
| | | DMD | 26.4 | 0.3 |
| | D25 | Healthy | 22.8 | 0.4 |
| | | DMD | 25.1 | 2.5 |
| ***Dp412e*** | D0 | Healthy | 37.2 | 1.7 |
| | | DMD | 35.8 | 2.4 |
| | D3 | Healthy | 33.3 | 1.9 |
| | | DMD | 32.7 | 2.1 |
| ***Dp427c*** | D0 | Healthy | 31.6 | 0.8 |
| | | DMD | 33.3 | 0.8 |
| | D3 | Healthy | 34.1 | 0.2 |
| | | DMD | 38.2 | 3.8 |
| | D10 | Healthy | 33.5 | 0.9 |
| | | DMD | 34.8 | 0.4 |
| | D17 | Healthy | 30.1 | 1.0 |
| | | DMD | 30.8 | 0.4 |
| | D25 | Healthy | 30.1 | 0.3 |
| | | DMD | 30.0 | 0.3 |
| ***Dp427m*** | D3 | Healthy | 35.2 | 0.9 |
| | | DMD | 34.7 | 0.9 |
| | D10 | Healthy | 28.5 | 1.2 |
| | | DMD | 29.4 | 0.8 |
| | D17 | Healthy | 22.5 | 1.1 |
| | | DMD | 23.6 | 0.1 |
| | D25 | Healthy | 20.4 | 0.4 |
| | | DMD | 21.9 | 1.4 |

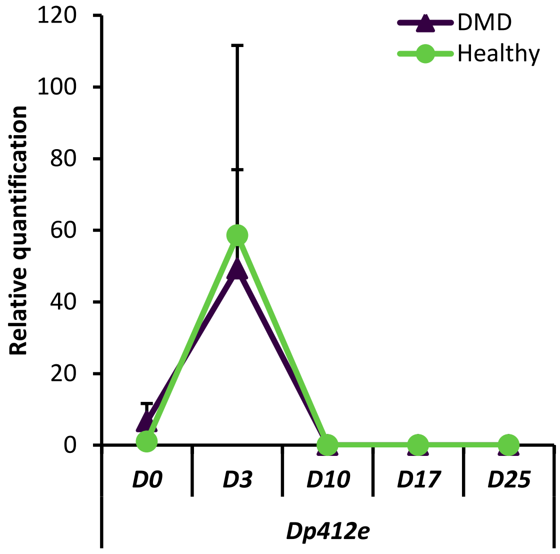


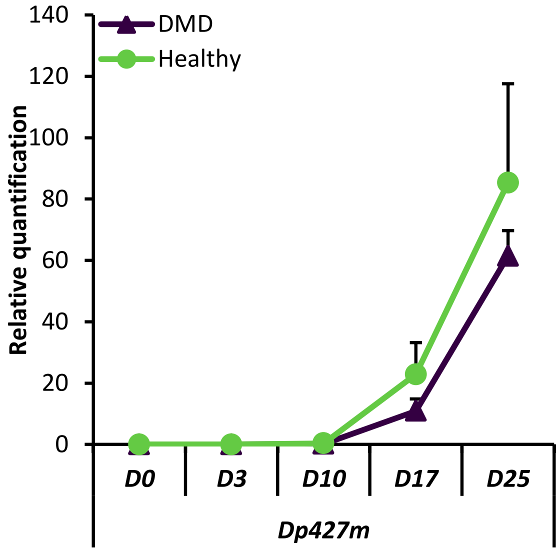


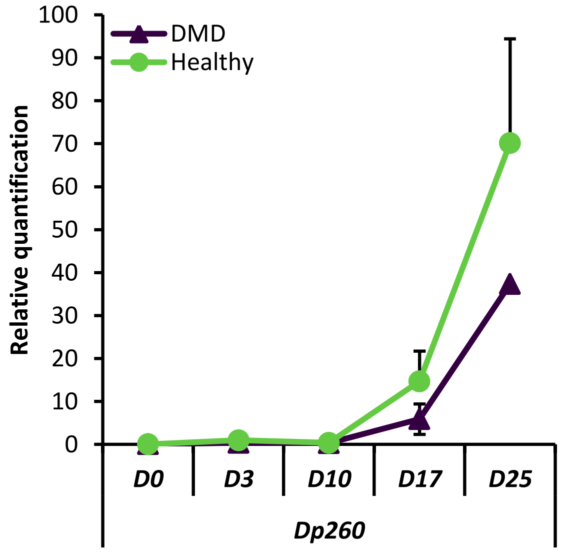

### Figure S2

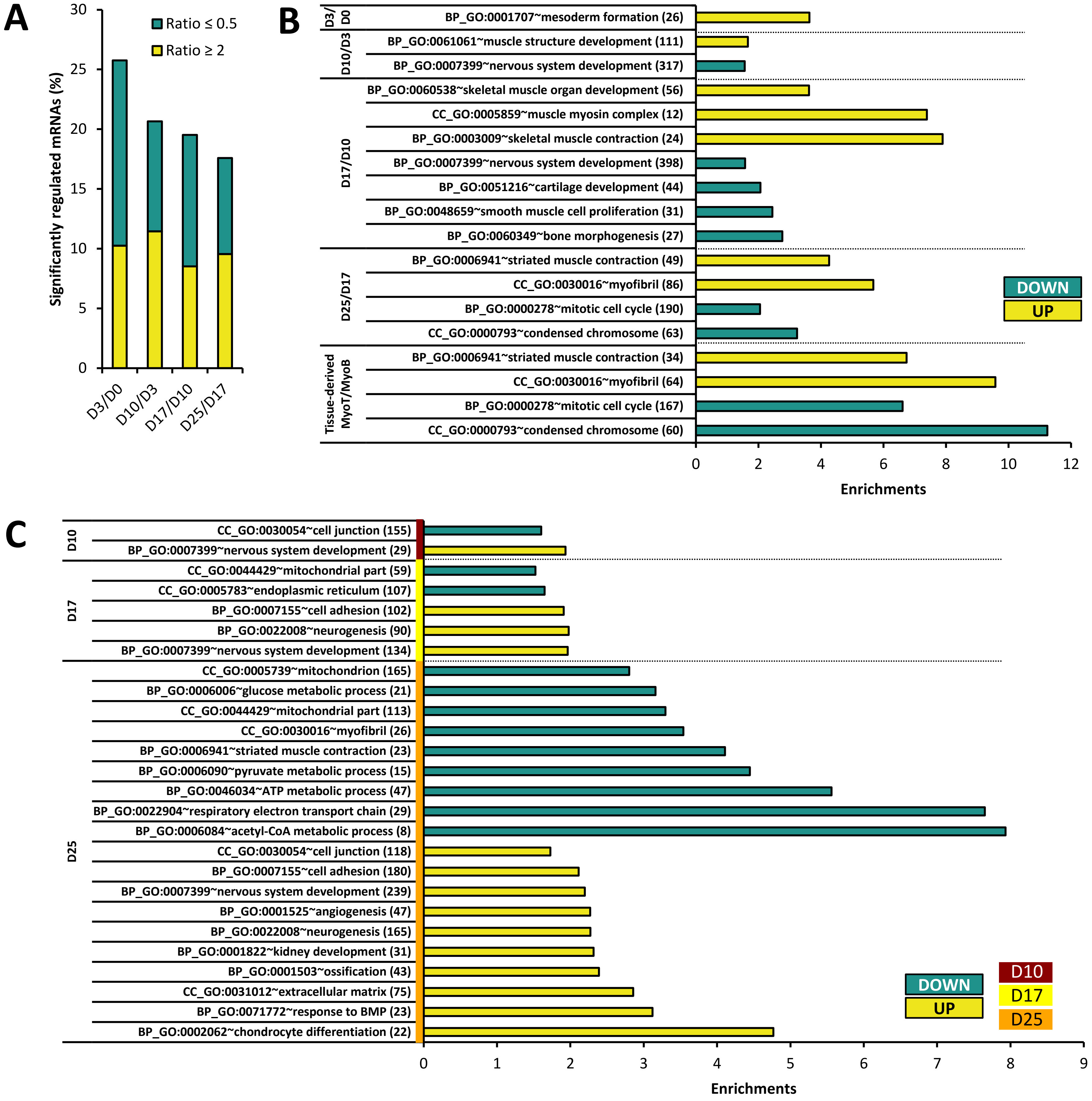
A
Significantly regulated mRNAs (%)
0
5
10
15
20
25
30
Ratio ≤ 0.5
Ratio ≥ 2
D3/D0
D10/D3
D17/D10
D25/D17
B
D3/D0
BP_GO:0001707~mesoderm formation (26)
D10/D3
BP_GO:0061061~muscle structure development (111)
BP_GO:0007399~nervous system development (317)
D17/D10
BP_GO:0060538~skeletal muscle organ development (56)
CC_GO:0005859~muscle myosin complex (12)
BP_GO:0003009~skeletal muscle contraction (24)
BP_GO:0007399~nervous system development (398)
BP_GO:0051216~cartilage development (44)
BP_GO:0048659~smooth muscle cell proliferation (31)
BP_GO:0060349~bone morphogenesis (27)
D25/D17
BP_GO:0006941~striated muscle contraction (49)
CC_GO:0030016~myofibril (86)
BP_GO:0000278~mitotic cell cycle (190)
CC_GO:0000793~condensed chromosome (63)
Tissue-derived MyoT/MyoB
BP_GO:0006941~striated muscle contraction (34)
CC_GO:0030016~myofibril (64)
BP_GO:0000278~mitotic cell cycle (167)
CC_GO:0000793~condensed chromosome (60)
0
2
4
6
8
10
12
Enrichments
DOWN
UP
C
D10
CC_GO:0030054~cell junction (155)
BP_GO:0007399~nervous system development (29)
D17
CC_GO:0044429~mitochondrial part (59)
CC_GO:0005783~endoplasmic reticulum (107)
BP_GO:0007155~cell adhesion (102)
BP_GO:0022008~neurogenesis (90)
BP_GO:0007399~nervous system development (134)
D25
CC_GO:0005739~mitochondrion (165)
BP_GO:0006006~glucose metabolic process (21)
CC_GO:0044429~mitochondrial part (113)
CC_GO:0030016~myofibril (26)
BP_GO:0006941~striated muscle contraction (23)
BP_GO:0006090~pyruvate metabolic process (15)
BP_GO:0046034~ATP metabolic process (47)
BP_GO:0022904~respiratory electron transport chain (29)
BP_GO:0006084~acetyl-CoA metabolic process (8)
CC_GO:0030054~cell junction (118)
BP_GO:0007155~cell adhesion (180)
BP_GO:0007399~nervous system development (239)
BP_GO:0001525~angiogenesis (47)
BP_GO:0022008~neurogenesis (165)
BP_GO:0001822~kidney development (31)
BP_GO:0001503~ossification (43)
CC_GO:0031012~extracellular matrix (75)
BP_GO:0071772~response to BMP (23)
BP_GO:0002062~chondrocyte differentiation (22)
0
1
2
3
4
5
6
7
8
9
Enrichments
DOWN
UP
D10
D17
D25

### Figure S3

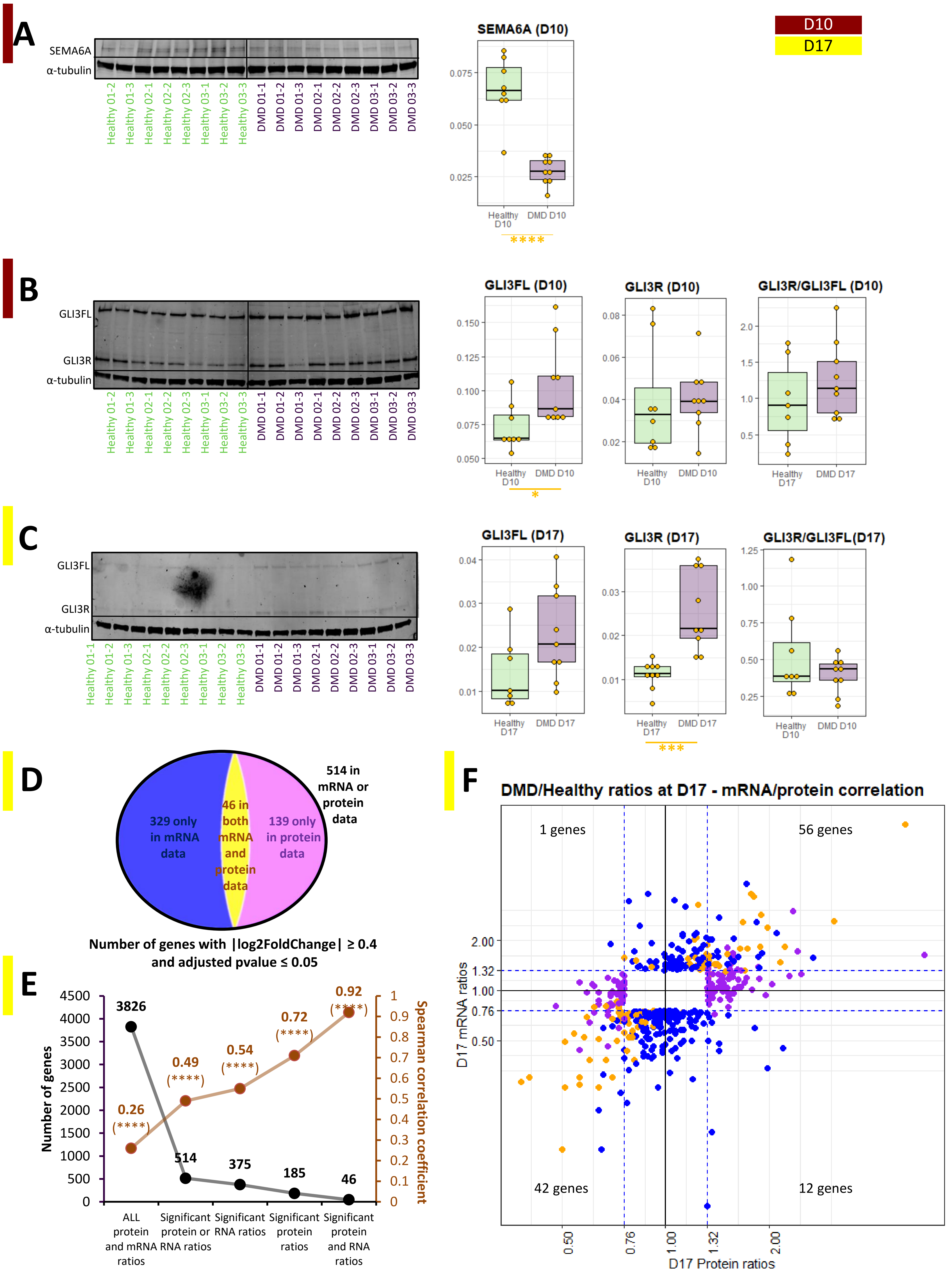

A
SEMA6A
α-tubulin
Healthy 01-2
Healthy 01-3
Healthy 02-1
Healthy 02-2
Healthy 02-3
Healthy 03-1
Healthy 03-2
Healthy 03-3
DMD 01-1
DMD 01-2
DMD 01-3
DMD 02-1
DMD 02-2
DMD 02-3
DMD 03-1
DMD 03-2
DMD 03-3
SEMA6A (D10)
Healthy D10
DMD D10
****
D10
D17
B
GLI3FL
GLI3R
α-tubulin
GLI3FL (D10)
GLI3R (D10)
GLI3R/GLI3FL (D10)
Healthy D17
DMD D17
*
C
GLI3FL (D17)
GLI3R (D17)
GLI3R/GLI3FL(D17)
Healthy 01-1
***
D
514 in mRNA or protein data
329 only in mRNA data
46 in both mRNA and protein data
139 only in protein data
Number of genes with |log2FoldChange| ≥ 0.4 and adjusted pvalue ≤ 0.05
E
Number of genes
Spearman correlation coefficient
3826
514
375
185
46
0.26 (****)
0.49 (****)
0.54 (****)
0.72 (****)
0.92 (****)
ALL protein and mRNA ratios
Significant protein or RNA ratios
Significant RNA ratios
Significant protein ratios
Significant protein and RNA ratios
F
DMD/Healthy ratios at D17 - mRNA/protein correlation
1 genes
56 genes
42 genes
12 genes
D17 mRNA ratios
D17 Protein ratios

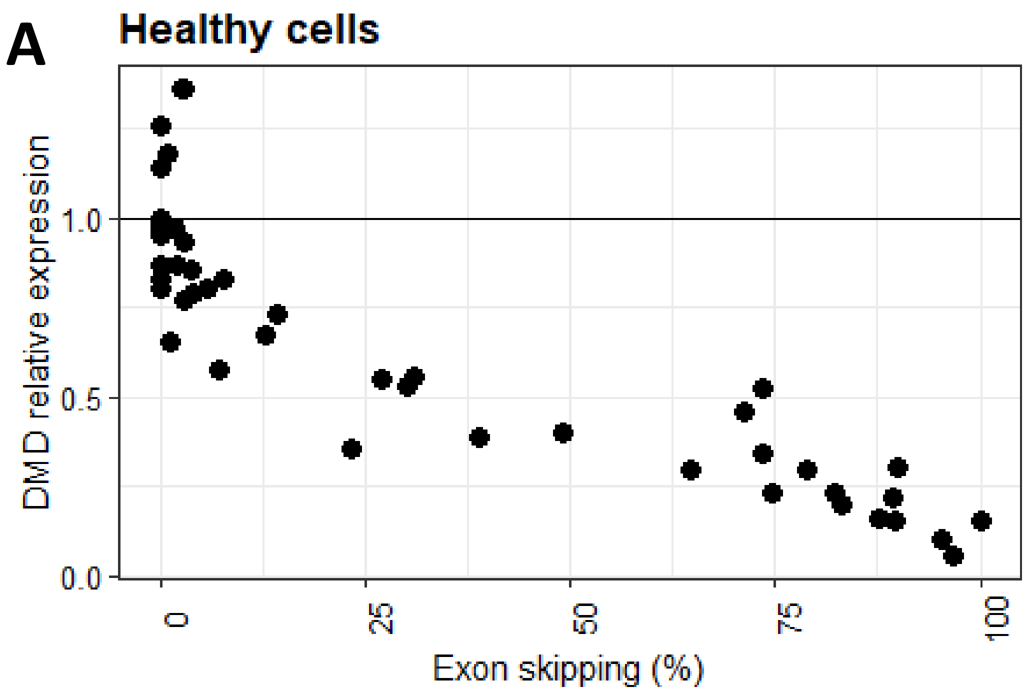


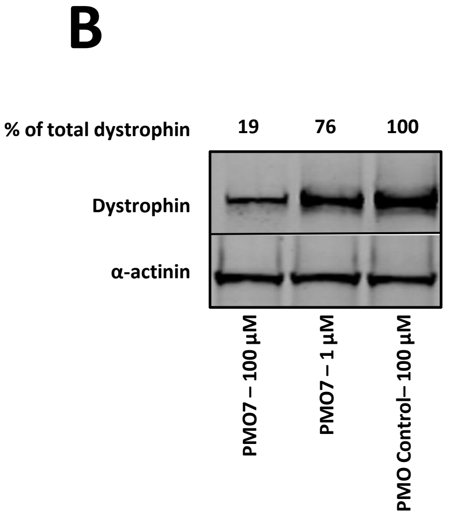


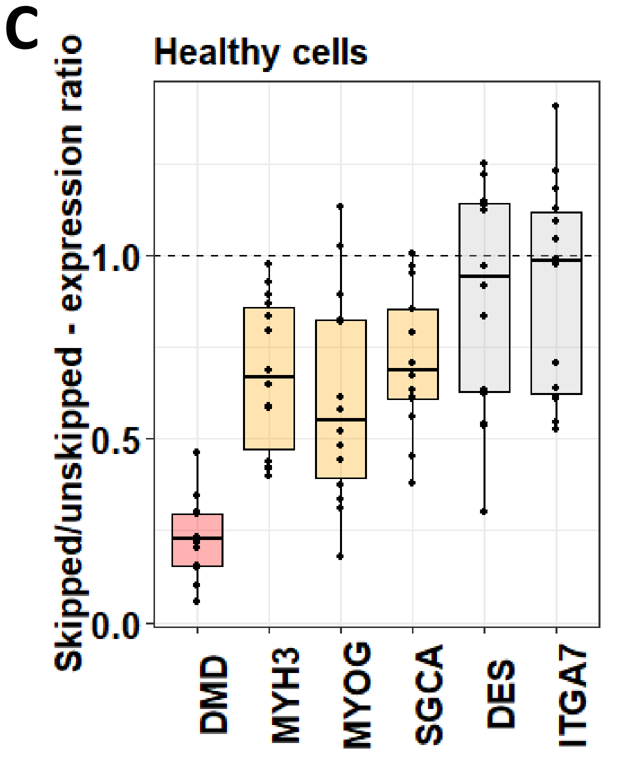


| | MYH3 | MYOG | SGCA | DES | ITGA7 |
|---|---|---|---|---|---|
| P value | **** | **** | **** | ns | ns |
| Spearman r | 0.56 | 0.64 | 0.5 | 0.15 | 0.062 |
| Number of analysed pairs | 59 | 59 | 56 | 59 | 59 |
| | Group 1 | | | Group 2 | |

### Figure S4

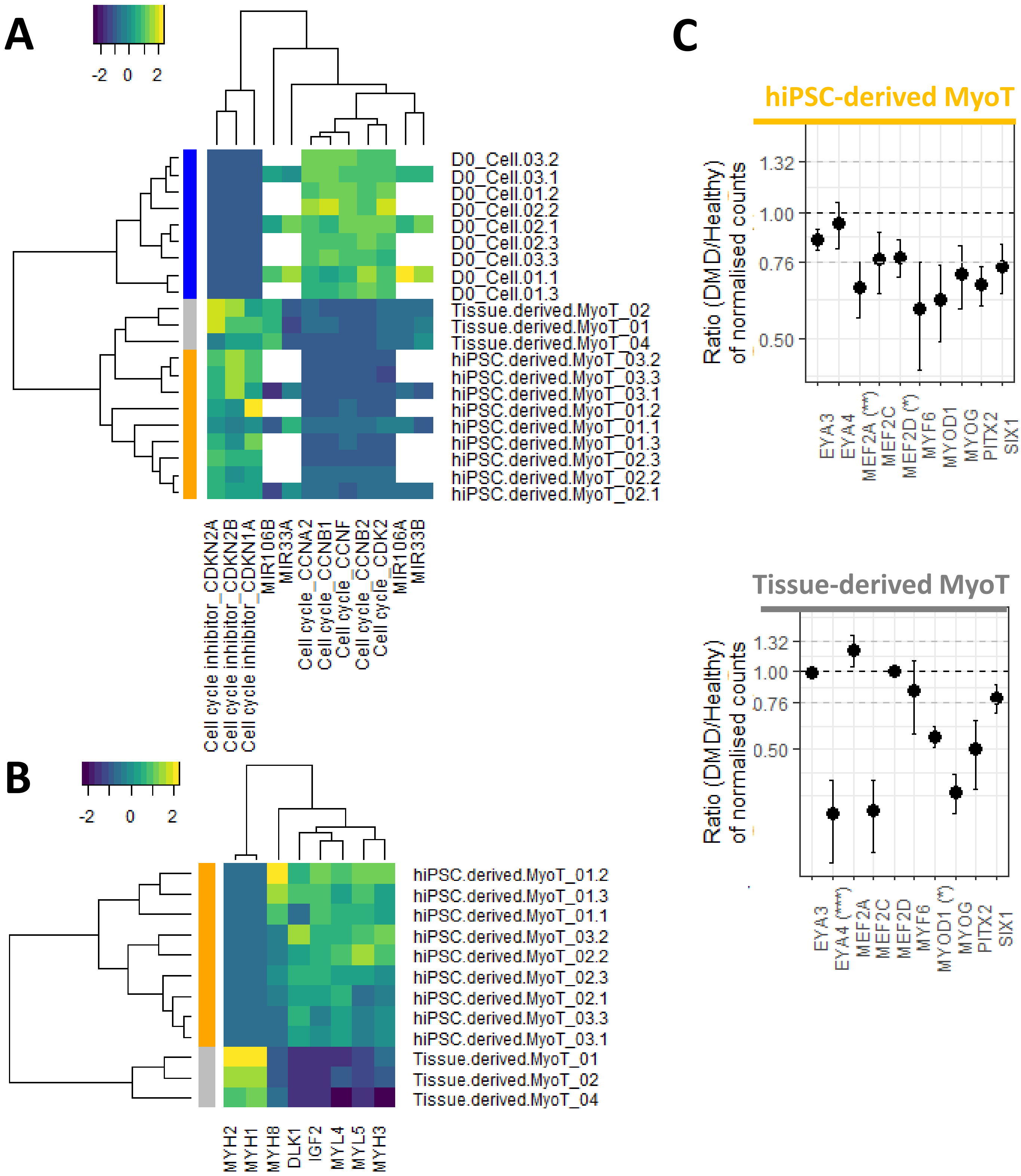
A
-2 0 2
D0_Cell.03.2
D0_Cell.03.1
D0_Cell.01.2
D0_Cell.02.2
D0_Cell.02.1
D0_Cell.02.3
D0_Cell.03.3
D0_Cell.01.1
D0_Cell.01.3
Tissue.derived.MyoT_02
Tissue.derived.MyoT_01
Tissue.derived.MyoT_04
hiPSC.derived.MyoT_03.2
hiPSC.derived.MyoT_03.3
hiPSC.derived.MyoT_03.1
hiPSC.derived.MyoT_01.2
hiPSC.derived.MyoT_01.1
hiPSC.derived.MyoT_01.3
hiPSC.derived.MyoT_02.3
hiPSC.derived.MyoT_02.2
hiPSC.derived.MyoT_02.1
Cell cycle inhibitor_CDKN2A
Cell cycle inhibitor_CDKN2B
Cell cycle inhibitor_CDKN1A
MIR106B
MIR33A
Cell cycle_CCNA2
Cell cycle_CCNB1
Cell cycle_CCNF
Cell cycle_CCNB2
Cell cycle_CDK2
MIR106A
MIR33B
B
-2 0 2
hiPSC.derived.MyoT_01.2
hiPSC.derived.MyoT_01.3
hiPSC.derived.MyoT_01.1
hiPSC.derived.MyoT_03.2
hiPSC.derived.MyoT_02.2
hiPSC.derived.MyoT_02.3
hiPSC.derived.MyoT_02.1
hiPSC.derived.MyoT_03.3
hiPSC.derived.MyoT_03.1
Tissue.derived.MyoT_01
Tissue.derived.MyoT_02
Tissue.derived.MyoT_04
MYH2
MYH1
MYH8
DLK1
IGF2
MYL4
MYL5
MYH3
C
hiPSC-derived MyoT
Ratio (DMD/Healthy) of normalised counts
1.32
1.00
0.76
0.50
EYA3
EYA4
MEF2A (**)
MEF2C
MEF2D (*)
MYF6
MYOD1
MYOG
PITX2
SIX1
Tissue-derived MyoT
Ratio (DMD/Healthy) of normalised counts
1.32
1.00
0.76
0.50
EYA3
EYA4 (****)
MEF2A
MEF2C
MEF2D
MYF6
MYOD1 (*)
MYOG
PITX2
SIX1

### Figure S5

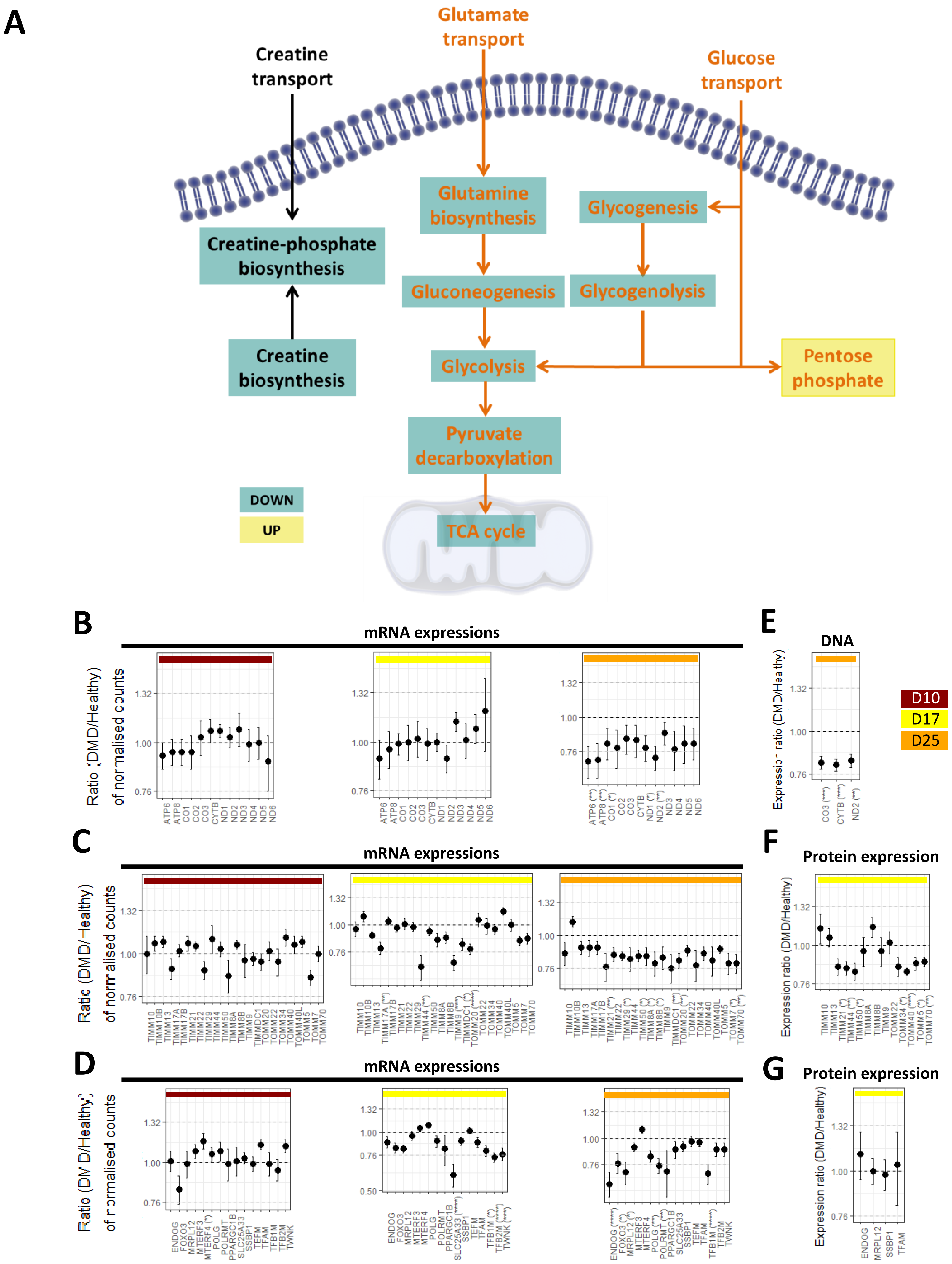
A
Creatine transport
Glutamate transport
Glucose transport
Creatine-phosphate biosynthesis
Glutamine biosynthesis
Glycogenesis
Gluconeogenesis
Glycogenolysis
Creatine biosynthesis
Glycolysis
Pentose phosphate
Pyruvate decarboxylation
TCA cycle
DOWN
UP
B
mRNA expressions
Ratio (DMD/Healthy) of normalised counts
C
mRNA expressions
D
mRNA expressions
E
DNA
Expression ratio (DMD/Healthy)
D10
D17
D25
F
Protein expression
G
Protein expression
